# The effect of recent habitat change on genetic diversity at putatively adaptive and neutral loci in *Primula veris* in semi-natural grasslands

**DOI:** 10.1101/2021.05.12.442254

**Authors:** S. Träger, C. Rellstab, I. Reinula, N. Zemp, A. Helm, R. Holderegger, T. Aavik

## Abstract

Recent habitat change in semi-natural grasslands due to a lack of management has been shown to affect the genetic diversity of grassland plants. However, it is unknown how a change in local environment affects genetic diversity at adaptive loci. We applied RADseq (restriction-site associated DNA sequencing) to extract > 3,000 SNPs across 568 individuals from 32 Estonian populations of *Primula veris*, a plant species common to semi-natural grasslands. We evaluated the effect of recent grassland overgrowth following management abandonment on the genetic diversity at putatively neutral and adaptive loci, which we distinguished by applying three methods, i.e., linear and categorical environmental association analyses, and an *F*_ST_ outlier test. For validation, we randomised the genotype to sample assignments. Effects of recent habitat change on genetic diversity differed between neutral and adaptive SNP sets. Genetic diversity assessed at putatively neutral loci was similar in open and overgrown habitats but showed a significant difference between these habitat types at putatively adaptive loci: overgrown (i.e. newly established) habitats exhibited higher genetic diversity at putatively adaptive loci than open (i.e. old) habitats, likely due to the exertion of novel selection pressures imposed by new habitat conditions. This increase in genetic diversity at putatively adaptive loci in the new environment points to currently ongoing selection processes where genetic adaptation to the old habitat is potentially lost through altered allele frequencies. Our study suggests that a recent change in local habitat conditions may not be reflected in neutral loci whereas putatively adaptive loci can inform about potential selection processes.

## Introduction

Habitat change is a consequence of ongoing anthropogenic landscape and climatic change (IPBES, 2018; IPCC, 2019). In particular, intensification of land use and abandonment of traditional management practices during the last century led to a dramatic degradation and isolation of habitats such as European semi-natural grasslands (Cousins, Auffret, Lindgren, & Tränk, 2015; Hooftman & Bullock, 2012). Historically, moderate and continuous human management (e.g. grazing by livestock) has supported development of open semi-natural grassland habitats with increased light availability and elevated niche partitioning, facilitating uniquely high levels of biodiversity (Habel et al., 2013; Wilson, Peet, Dengler, & Pärtel, 2012). Yet, conversion to other land use types or abandonment of traditional management have gradually caused substantial loss in grassland area and increased fragmentation, i.e. due to overgrowth with dense woody vegetation. This has resulted in negative effects on grassland biodiversity (Habel et al., 2013; Picó & Van Groenendael, 2007).

Likewise, the intra-specific genetic diversity of grassland plants suffers from the degradation, fragmentation, and loss of semi-natural grasslands. Due to potentially reduced population sizes and increased landscape barriers, the genetic diversity of many insect-pollinated grassland plant species is compromised by interrupted pollen-mediated gene flow (e.g. DiLeo, Holderegger, & Wagner, 2018; Tewksbury et al., 2002). This likely aggravated gene flow contributes to potentially decreased genetic diversity and increased genetic differentiation in plant populations in affected grasslands (e.g. Honnay & Jacquemyn, 2007; Picó & Van Groenendael, 2007). However, plants might exhibit a delayed response to habitat change (e.g. Aavik et al., 2019; Helm, Hanski, & Pärtel, 2006; Lehtilä et al., 2016). For instance, life history traits, such as mating system and lifespan, determine the speed and magnitude of a plant’s response to a changed environment (Hamrick & Godt, 1996; Leimu, Mutikainen, Koricheva, & Fischer, 2006). With many grassland plants having a relatively long lifespan of up to several decades (Ehrlén & Lehtilä, 2002), such longevity might mask genetic effects induced by habitat change.

Genetic diversity is one of the central parameters in estimating a population’s adaptive potential (Bilska & Szczecińska, 2016). However, because environmental change does not leave a signature in all parts of the genome (Nei, Suzuki, & Nozawa, 2010), such estimation requires differentiating between genetic diversity assessed at putatively neutral and adaptive loci (Bilska & Szczecińska, 2016). Here, we refer to putatively neutral loci when loci are affected by neutral processes such as gene flow but not by specific environmental factors. In contrast, putatively adaptive loci explicitly show a response to tested environmental factors. Well-adapted populations might show a reduced diversity at adaptive loci due to beneficial mutations going towards fixation, and it is the adaptive potential at the population level that can be assessed with investigations of genetic diversity at adaptive loci (Milot et al., 2020). However, the difference in response of genetic diversity at putatively neutral and adaptive loci to habitat change has been mostly ignored so far, in particular in the context of recent land use change. The only existing studies that explicitly account for differences in neutral and adaptive loci in plants concentrated on climatic factors (Dauphin et al., 2020; Sun et al., 2020). Moreover, in conservation genetics, most studies focussed on overall or genetic diversity at neutral loci, often using a set of neutral microsatellite markers, while ignoring adaptive regions of the genome (González et al., 2019; Wei & Jiang, 2020). However, it is exactly the adaptive regions that are important for the fate of a population.

With gradual grassland overgrowth, plant populations experience changed environmental conditions with novel selection pressures such as lower light availability, changes in soil chemical conditions, and altered pollinator communities (Helm, 2019), demanding for phenotypic plasticity of individuals or adaptation to the new habitat conditions. The adaptive potential of a population can be nourished from three sources: standing genetic variation (Barrett & Schluter, 2008; Radwan & Babik, 2012), gene flow (Slatkin, 1985) or, in the longer term, spontaneous mutations. Newly introduced barriers to gene flow should increase the importance of a population’s standing genetic variation, and novel selection pressures induced by habitat change likely trigger a decrease in adaptation to the former grassland habitat. Thus, habitat change forces plants to either adapt to locally new habitat conditions predominantly based on their standing genetic variation in an aggravated gene flow scenario, to disperse to more favourable habitats, or to face local extinction (e.g. Cheptou, Hargreaves, Bonte, & Jacquemyn, 2017; Frankham, 2005).

In the present study, we were interested in the effect of recent overgrowth of Estonian semi-natural grasslands with woody vegetation over the past century on the genetic diversity at putatively neutral and adaptive loci in *Primula veris* populations, a long-lived grassland specialist plant. Our study is one of the first to test for a land use change effect on both putatively neutral and adaptive loci in *in-situ* wild plant populations. We applied double-digest restriction-site associated DNA sequencing (ddRADseq) in 32 populations of *P. veris* from open and recently overgrown grasslands. We distinguished between putatively neutral and adaptive loci by performing a combination of environmental association analyses and *F*_ST_ outlier tests. We specifically asked whether (1) genetic diversity at neutral loci of *P. veris* populations is negatively affected by overgrowth of grasslands due to potentially reduced population size and/or aggravated gene flow; (2) genetic diversity at adaptive loci exhibits a different response to habitat change than genetic diversity at neutral loci; and (3) genetic diversity at adaptive loci is actually increased due to ongoing selection processes where genetic adaptations to the old, open habitat are slowly lost through altered allele frequencies of the beneficial alleles for open and overgrown habitats.

## Materials and Methods

### Study species

*Primula veris* L. (Primulaceae) is an herbaceous perennial rosette-forming hemicryptophyte most commonly occurring in calcareous grasslands. *Primula veris* prefers open habitats but can grow under shade with reduced reproduction (Brys & Jacquemyn, 2009). Its average life span reaches up to 50 years (Ehrlén & Lehtilä, 2002). In Estonia, the study region, *P. veris* generally flowers in May. The study species is an obligate outbreeder that depends on insect-pollination (Deschepper, Brys, & Jacquemyn, 2018). Pollen dispersal is spatially restricted to several meters (Brys & Jacquemyn, 2009). Self-pollination is prevented by heterostyly with two flower morphs, with low levels of successful intra-morph pollination (Wedderburn & Richards, 1990). Primary seed dispersal is limited to a few metres from the maternal plant (Brys & Jacquemyn, 2009).

### Study sites and sampling

Study sites were located on dry calcareous grasslands, alvars, on the islands of Muhu and Saaremaa in Western Estonia (Figure 1). Alvars are semi-natural grasslands on Ordovician and Silurian bedrock with only a low soil depth (< 20 cm). Management, i.e. grazing livestock, in the area was abandoned 20 – 90 years ago. Our study sites were part of a large-scale biodiversity inventory of the European Commission’s LIFE+ Nature program restoration project “ LIFE to Alvars” (Helm, 2019), which included monitoring of genetic diversity of grassland plant species, including *P. veris*, in alvars at different successional stages of overgrowth (e.g. still open and recently overgrown). The mean temperature in the area is 17°C in summer and -3°C in winter, and the mean annual precipitation is about 680 mm (EWS, 2020).

**Figure 1.**
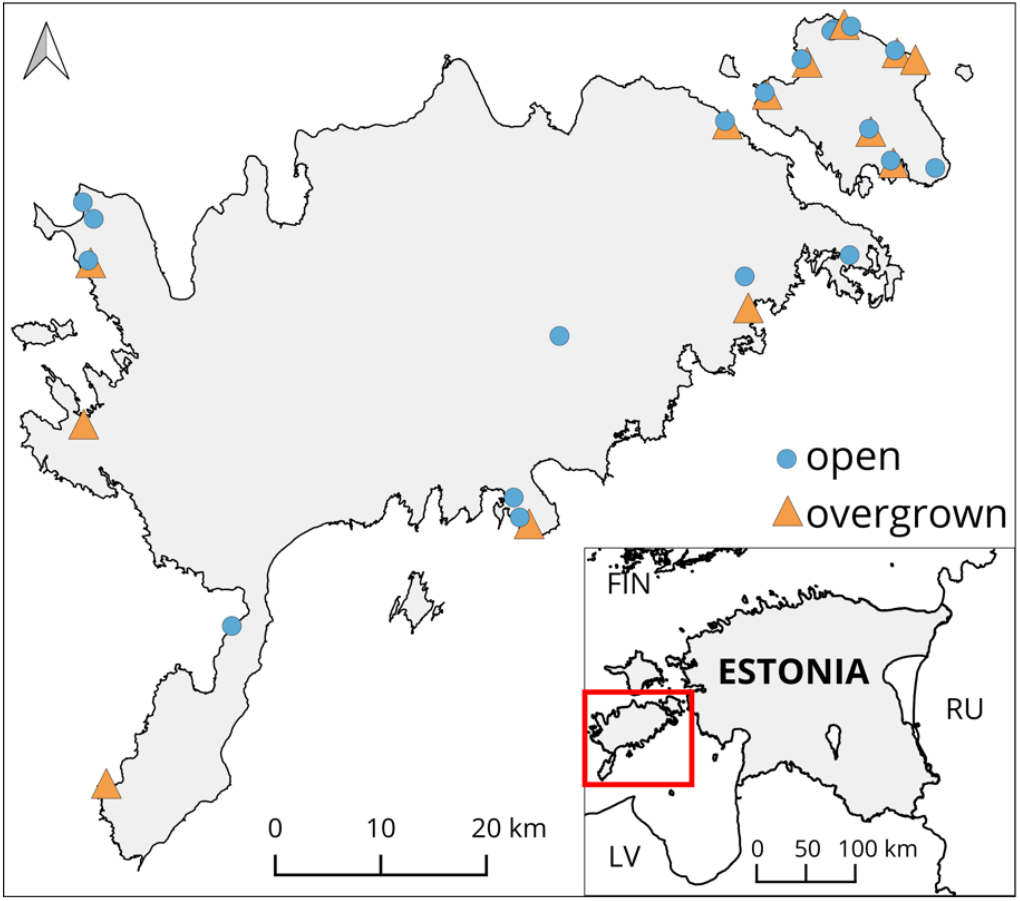
Map of the study area on Muhu and Saaremaa islands in Western Estonia. *Primula veris* populations are depicted with blue filled circles when occurring in open semi-natural grassland and with orange filled triangles when occurring in recently overgrown historical semi-natural grasslands.

We sampled 32 populations (i.e. spatially distinct patches) of *P. veris* distributed across two regions, Muhu and Saaremaa islands, in the summers of 2015 and 2016 (Figure 1; Table 1). Where possible, we chose pairs of closely located populations (i.e. within pollen- and seed-mediated gene flow distance) of contrasting habitat types (i.e. open and recently overgrown grasslands). Nineteen populations were located in open grasslands (i.e. old habitat; hereafter open habitats) and 13 populations were located in shrubby-overgrown grasslands (i.e. new habitat; hereafter overgrown habitats), comprising 10 population pairs with an average distance of 533 m and a minimum distance of 20 m between members of pairs. Such a paired sampling design has been shown to be efficient in detecting genomic signatures of local adaptation in environmental association analysis (EAA; Lotterhos & Whitlock, 2015) and allows the use of categorical EAA approaches (see below). Overgrown habitats represented mid-successional stages with at least 60% cover of shrubby vegetation, mostly *Juniperus communis*.

**Table 1.**
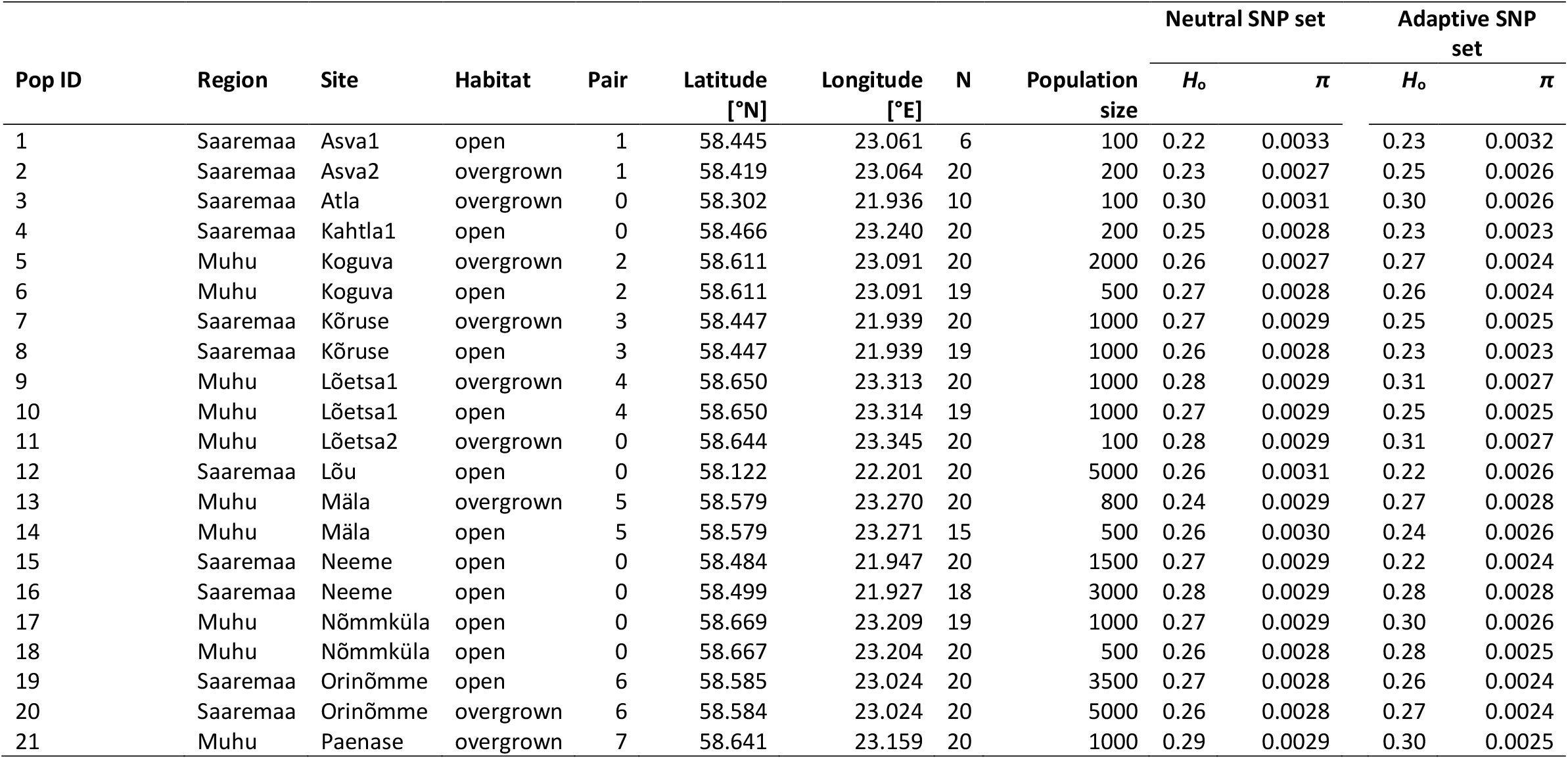
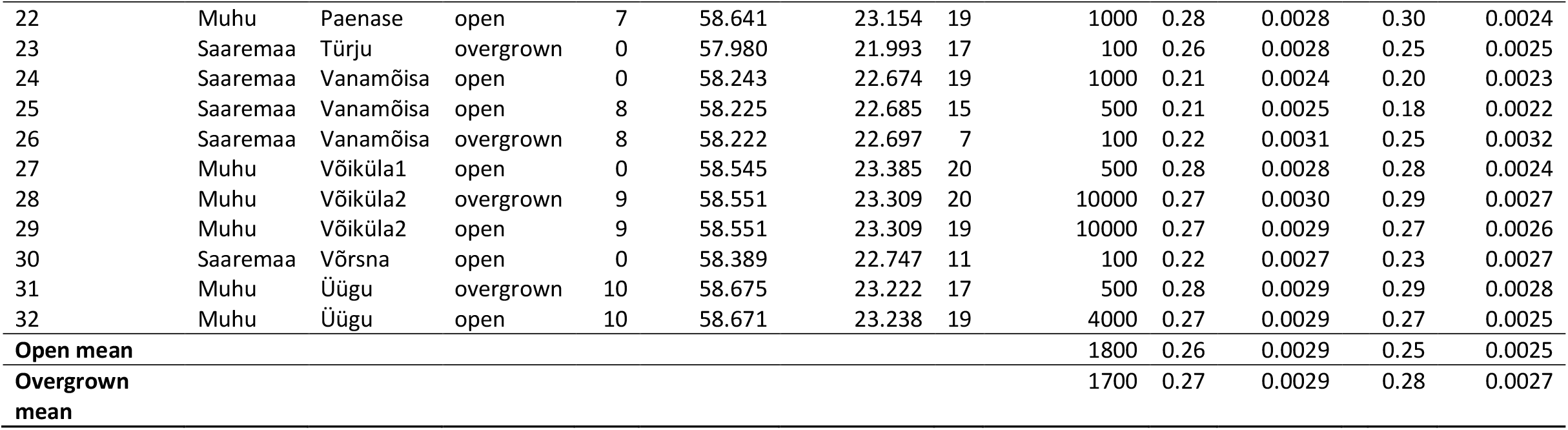
Location information and genetic diversity measurements for the studied populations of *Primula veris* in Muhu and Saaremaa, Estonia. Habitat – affiliation to open or overgrown grassland; pair – affiliation to a pair of closely situated populations (0 indicates affiliation to no pair); N – sample size after SNP filtering; population size – estimated number of individuals per population; *H*_o_ – observed heterozygosity; *π* – nucleotide diversity.

Within each population, we sampled three fresh leaves of 20 random flowering *P. veris* individuals (where possible) that were at least 50 cm apart. Leaves were stored in silica-gel until further processing. Approximate population census sizes of *P. veris* were estimated by assessing the number of both flowering and non-flowering individuals per population.

### Environmental data

To characterize the environment of the study sites, we used data collected within the frame of the “ LIFE to alvars” project (Helm, 2019). We selected 16 *in-situ* measured environmental variables with the potential to represent contrasting habitat types (open and overgrown), i.e. “ openness”, “soil”, and “biota”. For openness, we considered the total percentual shrub and tree coverage, respectively, assessed within a radius of 10 m from the center of *P. veris* populations, and the light availability above and below the herbal layer measured with a Li-Cor LI-250 Light Meter and a LI-190SA Quantum Sensor (Lincoln, Nebraska, USA), in 1×1 m in the center of the population. For soil, we considered average soil depth in cm based on ten random locations taken within a 10 m radius around the central point of *P. veris* populations. In the same radius, five soil samples were taken from random locations and pooled for chemical analyses. From each sample, soil pH (KCl solution), available soil phosphorus (P; extraction with acid ammonium lactate solution), potassium (K), magnesium (Mg), calcium (Ca), and soil organic content (OC, loss on ignition) were measured. For biota, we considered butterfly and bumblebee abundance and richness and vascular plant richness within a 10 m radius around the central points of *P. veris* populations. Butterflies and bumblebees were monitored using standardised transect counts (Pollard, 1977). Each site was visited three times for butterflies and two times for bumblebees over two years to cover phenological aspects of different species (Helm, 2019).

In addition to the 16 *in-situ* measured environmental variables, we extracted climate data for each population from CHELSA (Karger et al., 2017) with a resolution of 30 arc sec from the reference period 1979-2013. To increase resolution, we applied a bi-linear interpolation that accounts for the climate values in surrounding grid cells and the position of the population within the grid cell. For our analyses, we used temperature (Bio1, annual mean temperature) and precipitation (Bio12, annual precipitation sum), because they represent the most comprehensive bioclimatic variables describing the climate in our study region.

To test which environmental factors significantly differed between the two habitat types, we performed a (non-paired) two-sample t-test for each of the 18 environmental variables.

### DNA extraction and ddRAD sequencing

Twenty-five mg of leaf material were pulverized with 2.3-mm chrome-steel beads (BioSpec Products, Bartlesville, USA) in a Mixer Mill 301 (Retsch, Haan, Germany). DNA was extracted using the LGC sbeadex plant kit (LGC, Berlin, Germany). 400 µl lysis buffer, consisting of 1% RNAse (100 mg/ml), 0.2% Proteinase K solution (20 mg/ml), and lysis buffer PN were added to pulverized samples, with an incubation time of 1 h at 65°C on a Thermomixer comfort (Eppendorf, Hamburg, Germany) at 300 rpm, and followed by a centrifugation at 2500 x g for 10 min. Lysates were transferred to binding solution, consisting of 420 µl binding buffer PN and 10 µl sbeadex particle solution. All following steps were conducted on a KingFisher Flex Purification System (Thermo Fisher Scientific, Waltham, USA), with the specification of using 400 µl of wash buffer PN1 twice, 400 µl of wash buffer PN2, and eluting purified DNA in 50 µl elution buffer AMP.

We applied a ddRADseq procedure by customizing an existing ddRADseq protocol (Westergaard et al., 2019). ddRADseq applies a double restriction enzyme digest followed by a size-selection of genomic fragments (Peterson, Weber, Kay, Fisher, & Hoekstra, 2012). RADseq provides a simple and cost-effective method to uncover thousands of polymorphic markers, both neutral and adaptive, in model and non-model organisms (e.g. Davey et al., 2011). Because the aim of our study was to identify general patterns of genetic diversity assessed at neutral and adaptive loci across a high number of samples from many populations, rather than identifying specific genes involved in adaptation, we chose not to use whole genome or targeted sequencing. Such methods might be, however, worthwhile to consider in future analyses following the results of the present study.

For the detailed ddRADseq protocol see supplemental information (Supplemental Methods). Briefly, standardized DNAs of fully randomized samples were digested and purified before ligation to a combination of one of 48 EcoRI and 2 TaqI adapters, respectively, resulting in uniquely tagged barcoded DNA samples. DNA samples with the same TaqI adapter but different EcoRI adapters (48 samples) were pooled together and size selected for fragments of 450 bp length. The size-selected sample pools were selected for fragments containing biotin labelled TaqI adapters. Subsequently, polymerase chain reaction (PCR) was conducted, PCR products (ddRADseq libraries) were purified and their DNA concentration was measured to calculate molarity per ddRADseq library. Finally, samples with distinct TaqI multiplexing indices were combined to produce a final library of at least 5 nM consisting of 96 samples (2 × 48 uniquely barcoded samples from two multiplex indices). In addition, for sequencing, we used 15% of a standard Illumina library to increase index diversity.

Pooled libraries were prepared according to guidelines of the sequencing facility and sequenced on an Illumina HiSeq2500 at the Functional Genomics Centre Zurich (FGCZ, Switzerland), using one lane per library with 125 cycles in single-end read (125 bp), high-output mode. The sample set per library included a negative (no sample DNA) and a positive (sample replica; different positive controls in different libraries) control to exclude the possibility of contamination and to calculate the genotyping error of SNPs.

### Bioinformatic analysis

Sequence data were demultiplexed, and PCR duplicates were filtered using the functions “ process_radtags” and “ clone_filter” of STACKS v1.47, respectively (Catchen et al., 2011; Catchen, Hohenlohe, Bassham, Amores, & Cresko, 2013). We used trimmed sequencing reads (generated sequences), applying TRIMMOMATIC v0.36 (Bolger, Lohse, & Usadel, 2014) with the following conditions: (1) removing Illumina adapter matches allowing a maximum of two mismatches, (2) removing leading and trailing low quality or N bases below a quality score of 5, (3) performing a 5 bp sliding window quality check and trimming sequence ends if quality dropped below 15, and (4) dropping sequences, which were < 50 bp after previous quality checks. Pre-filtered sequence reads were aligned and mapped against the reference genome of *P. veris* (Nowak et al., 2015) using BURROWS-WHEELER ALIGNER v0.7.17 (BWA; Li, 2013). SNPs were called using FREEBAYES v1.1.0-54-g49413aa (Garrison & Marth, 2012) applying default values except a minimum-mapping-quality of 5, a minimum-base-quality of 5, and evaluating the 10 best SNP alleles. We only used SNPs which met quality criteria of the DDOCENT SNP filtering pipeline (Puritz, Hollenbeck, & Gold, 2014; Puritz, Matz, et al., 2014) with customizing the following parameters: minimum quality score of 20, minor allele count of 3, and maximum missing value proportion of 20% across all individuals. Loci potentially in linkage disequilibrium were filtered using VCFTOOLS 0.1.15 (*geno-r2* function; Danecek et al., 2011) keeping one random SNP of potentially linked SNP pairs with a threshold of 0.8. Loci showing an excess of heterozygotes (> 60% of the samples identified as heterozygotes) were filtered in R v3.4.2 (R Development Core Team, 2017). The genotyping error of filtered SNPs was calculated by the weighted mean of error rates using replica samples (i.e. positive controls) with TIGER v1.0 (Wegmann Lab, 2019).

### Compilation of adaptive and neutral SNP sets

From the total SNP data set (SNP_overall), we identified putatively adaptive SNPs (SNP_adapt) that were either (a) linearly associated to environmental factors or (b) categorically associated to habitat types (see below). An additional putatively adaptive SNP set (SNP_BayeScan) was compiled by identifying SNPs under potential diversifying or balancing selection applying an *F*_ST_ outlier test using BAYESCAN v2.1 (Foll & Gaggiotti, 2008). The models for BAYESCAN ran with default parameters (10 prior odds, 5,000 iterations with a thinning interval of 10, a burn in of 50,000 and 20 pilot runs of 5,000 iterations). Potential *F*_ST_ outlier loci were extracted for *q*-values 0.05.

The putatively neutral SNP set (SNP_neutral) was gained by excluding both putatively adaptive SNP sets (SNP_adapt and SNP_BayeScan) from SNP_overall. Subsequently, we calculated population genetic diversity parameters for both SNP_neutral and SNP_adapt. We did not use the SNP_BayeScan set for the estimation of genetic diversity indices at adaptive loci, because the SNPs identified by BAYESCAN might represent loci putatively involved in adaptation to other environmental factors than investigated here and were only used to better define SNP_neutral.

### Environmental association analyses

To detect putative signatures of natural selection in open and overgrown habitats, we used EAA that correlates environmental variation (describing the local habitat) with genetic variation of a population (Rellstab, Gugerli, Eckert, Hancock & Holderegger, 2015). Here, we performed EAAs with two types of relationships: linear and categorical.

For the linear analysis, we used latent factor mixed models (*lfmm_ridge* and *lfmm_test* functions in LFMM v2.0 in R; Caye, Jumentier, Lepeule, & François, 2019), which test for a linear relationship of the allele frequency (AF) at each SNP with each environmental variable while accounting for population structure with random latent factors. All 32 populations were considered, and all 18 environmental variables were used in EAA. We did not remove correlated variables, because our aim was to identify all SNPs with any sign of environmental adaptation for the compilation of the different SNP sets. For all subsequent analyses, however, we concentrated on those SNPs that were associated with environmental variables that significantly differed among the two habitat types. In LFMM, the number of latent factors must be set by the user and is recommended to be based on the number of genetic clusters in the study systems (see below) and the inflation factor *λ* (François, Martins, Caye, & Schoville, 2016). To control for false discoveries, we adjusted the *p* values per environmental variable using *λ* and the *χ*^2^ distribution (Caye et al., 2019; François et al., 2016) and applied the Benjamini-Hochberg algorithm (Benjamini & Hochberg, 1995) with a false discovery rate (FDR) of 0.05.

For the categorical analysis, we performed three different pairwise analyses on population AFs of the 20 populations that were sampled in pairs, i.e., geographically close, but environmentally diverged (open-overgrown; Table 1): a paired t-test, a paired Wilcoxon signed-rank test, and a sign test. To reduce false positive findings, only SNPs whose population AFs were significantly different between habitat types in all three categorical tests were considered for further analyses. For the t- and Wilcoxon tests, we used an *α* value of 0.05. In the sign test, we checked whether AF differences between the two populations of all pairs were consistent (i.e. had the same sign). We considered SNPs significant if they were consistent in a minimum of eight out of ten comparisons (pairwise AF differences of 0 were treated as consistent). In these categorical analyses, population structure is not directly incorporated, but since they test for differences within pairs, population structure can be ignored, as it is unlikely that population structure would lead to different signs of AF differences in different pairs. For all further analyses on putatively adaptive SNPs (SNP_adapt), we concentrated on the SNPs that were (a) associated in LFMM to those environmental variables that significantly differed among the two habitat types and/or (b) were significant in all three categorical tests.

For the SNP_adapt set, we also wanted to know whether we find a non-random pattern of AF change (“direction”) between the two populations of habitat pairs. For each SNP, we identified the putatively beneficial allele (i.e. major allele; AF > 0.5) for the open habitat and then calculated the average AF change of this beneficial allele from open to overgrown habitat (the historical habitat change) within pairs. We then counted how many SNPs exhibited a beneficial AF decrease or increase from open to overgrown habitats and used an exact two-sided binomial test in R to check if this pattern deviated from a random 1:1 ratio.

The use of linear and categorical EAAs for extracting putatively adaptive loci in respect to specific environmental factors to test the effect of habitat differences (which are caused by the same environmental factors) on genetic diversity indices might result in a circularity problem. This is because major AF and heterozygosity are not completely independent (Rosenberg & Jakobsson, 2008) and thus, differences in genetic diversity between habitat types might reflect the results of EAAs. To address this potential problem, we created 100 SNP_overall data sets with fully randomised genotype to sample assignments in order to create an empirical null distribution of the calculated parameters and statistics. Each of those 100 SNP_overall data sets (hereafter randomised data sets) went through the same process of EAAs and *F*_*ST*_ outlier analysis as the original data set to extract putatively neutral and adaptive loci.

### Population genetic diversity

For measurements of genetic diversity at putatively neutral and adaptive loci, we calculated, for all SNP sets of the original and for SNP_neutral and SNP_adapt of each randomised data set, observed heterozygosity (*H*_*o*_) using GENALEX 6.503 (Peakall & Smouse, 2012), and mean nucleotide diversity (*π*) using VCFTOOLS within a window of 125 bp over all loci for each population.

The effect of habitat type (open or overgrown) and SNP set (SNP_neutral or SNP_adapt), and their interaction, on genetic indices (*H*_o_, *π*) was tested using linear mixed effect models with population (1 – 32) and region (Muhu or Saaremaa) as random effects (lmerTest v3.1-0, *lmer* function; Kuznetsova, Brockhoff, & Christensen, 2017), for the original and each randomised data set. The latter random effect accounted for potential differences in landscape history among regions. To quantify the importance of fixed factors and their interaction we used the Likelihood Ratio Test to obtain *p* values, analysing the variance between the full and reduced models with Satterthwaite approximation (*χ*^2^ and associated p values). Here, we were particularly interested in the interaction of habitat type and SNP set. A significant interaction implies that genetic diversity indices assessed at putatively neutral and adaptive loci behave differently in open and recently overgrown habitats. If the interaction term was found significant, we tested for differences of genetic diversity indices in the different habitats for each SNP set using post-hoc tests with least-square means (lsmeans v2.30-0, *lsmeans* function; Lenth, 2016). We also tested for an effect of population size on genetic diversity indices by including it as a fixed effect in the above mixed effect models. Due to non-significant effects of population size and its interaction, we only present the results of mixed effect models without population size.

We compared results of the original data set with those of the randomised data sets by (a) calculating a “permutation *p*-value” for each factor (habitat type, SNP set and their interaction), i.e. the number of *p*-values of the randomised data set model outputs which were smaller than the *p*-value of the original data set model output divided by the total amount of randomisations (100); (b) inspecting whether the proportion of putatively adaptive SNPs with decreasing and increasing beneficial AF, respectively, of the original data set is significantly different than the distribution of the proportion of SNP_adapt with decreasing and increasing beneficial AF of the randomised data sets, using a t-test; and (c) calculating the mean *H*_o_ and *π* per population over all 100 randomised data sets and conducting a mixed-effect model analysis as done for the original data set.

### Population genetic structure and potential gene flow

Population genetic structure using SNP_neutral and SNP_overall was analysed using discriminant analysis of principle components (DAPC) in adegenet v2.1.1 in R (Jombart, 2008). DAPC uses uncorrelated principal component analysis (PCA) variables for discriminant analysis, producing synthetic discriminant functions that maximize between-group variation while minimizing within-group variation (Jombart, Devillard, & Balloux, 2010). We used cross-validation with 50 replicates to determine the number of principal components (PCs) to be retained to avoid overfitting. The function *find*.*clusters* was used to determine the optimal number of clusters within the data sets. For validation, we also applied a hierarchical clustering tree analysis on a Nei’s genetic distance matrix using mmod v1.3.3 and the *aboot* function from poppr v2.8.4 in R (Kamvar et al., 2019), with a cut-off of 50 and a bootstrap sample of 1,000.

Pairwise genetic differentiation (*F*_ST_) among populations for SNP_neutral and SNP_overall was calculated using genepop v1.0.5 (Rousset et al., 2017) in R. Potential effects of geographic distance and habitat type “distance” (open-open, overgrown-overgrown, open-overgrown/overgrown-open) on genetic differentiation (*F*_ST_) were tested for both SNP sets using multivariate generalized linear mixed models fitted with Markov chain Monte Carlo techniques (MCMCglmm) with 2,000,000 iterations and 500,000 burnins (MCMCglmm 2.29, *MCMCglmm* function; Hadfield, 2010) to account for non-independence of pairwise distance data. We present results of the best model according to DIC for each SNP set.

After a first visual inspection of the genetic and geographic distance relationship, we fitted multiple simple linear functions (package stats 3.4.2, *lm* function; Chambers, 1992) to further characterize potential isolation by distance (IBD; Van Strien, Holderegger, & Van Heck, 2015) and to estimate the maximum geographic distance up to which gene flow as indicated by genetic differentiation might be prevalent compared to other genetic processes, such as genetic drift. One set of linear models included pairwise *F*_ST_ values as response variable and increasing geographic distance as explanatory variable, the other complementary set of linear models fitted *F*_ST_ against a constant for the difference in geographic distance to 100 km (i.e. the maximum distance between study populations). The threshold for potential gene flow was estimated as the point where the sum of the residual standard errors of sets of complementary models stayed constant.

## Results

Sequencing of ddRADseq fragments yielded on average about 150 M raw sequences per library, with on average about 1.2 M sequences per sample. SNP calling and quality filtering resulted in 4,588 SNPs. From those, 3,084 SNPs remained after excluding loci potentially in linkage equilibrium and with an excess of heterozygotes, in a total of 568 individuals from 32 populations. The genotyping error of quality-filtered SNPs was 0.004. Negative controls did not result in sequences.

### Putatively adaptive loci

From the 18 environmental variables describing the habitat of populations, six significantly (t-test, *p* ≤ 0.05) differed among the two habitat types (open and overgrown; Figure S 1): shrub coverage, light above and below the herbal layer, butterfly species richness and abundance, and plant species richness.

In the linear EAA, based on the number of clusters in the DAPC (*K* = 6, see below) and based on the fact that the inflation factor *λ* in un-adjusted *p* values did not vary considerably from *K* = 3-10 in all environmental variables, we chose *K* = 6 latent factors for the LFMM analysis. With an FDR of 0.05, we identified eight SNPs being associated to an environmental variable (Table S 1). Only three of them were associated to one of the six variables that significantly differed between the habitat types (butterfly abundance).

In the categorical EAA comparing the two habitat types, we identified 99 SNPs with the paired t-test (*p* ≤ 0.05), 95 with the Wilcoxon test (*p* ≤ 0.05), and 557 SNPs with the sign test. Seventy-four SNPs were identified in all three pairwise tests, but none of them overlapped with the three LFMM SNPs (Figure S 2). The 74 SNPs from categorical EAAs and the three SNPs from linear EAAs were used for further analyses (SNP_adapt = 77 SNPs). Using randomised data sets, the mean number of SNP_adapt was 82 (± 20 SD) SNPs.

In the 77 SNPs that were putatively involved in adaptation to habitat type, 53 SNPs showed a decrease of the average beneficial AF (for the old, open habitat) in the new, overgrown compared to the open habitat (Figure 2). The binomial test revealed that this pattern was significantly different from a random expectation (*p* < 0.01). However, AF differences between habitat types were small; average AF change was 0.09 (range 0.04-0.16) in the 53 SNPs with decreasing, and 0.08 (range 0.03-0.15) in the 24 SNPs with increasing beneficial AF. Yet, the maximum AF difference found between two populations of a pair in any SNP was 0.56 (Figure S 3). Using randomised data sets, the mean proportion of SNP_adapt loci with decreasing beneficial AF was 52 (± 9 SD) %, and the mean proportion of SNP_adapt loci with increasing beneficial AF was 48 (± 9 SD) % (Figure S 4).

**Figure 2.**
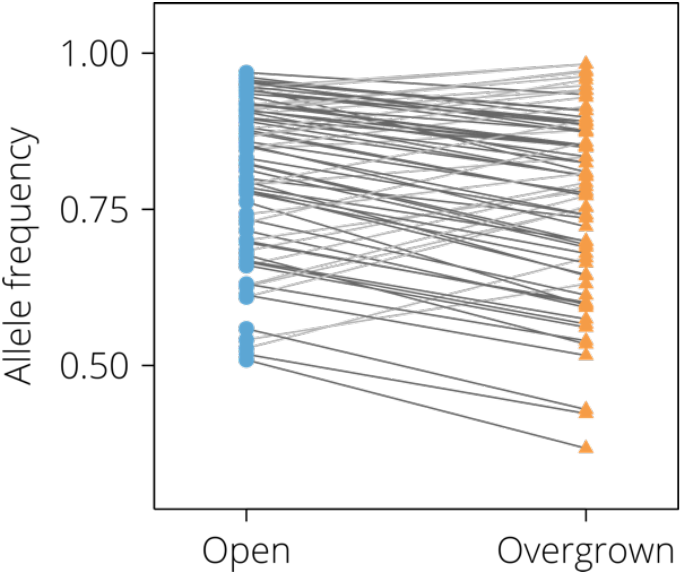
Patterns of allele frequency (AF) change of the 77 putatively adaptive SNPs (SNP_adapt) averaged across pairs of populations for each SNP. For each SNP, the putatively beneficial allele frequency for the open habitat (AF > 0.5) is shown. SNPs were derived from linear and categorical environmental association analyses. SNPs with decreasing/increasing AF in overgrown habitats (orange triangles) compared to open habitats (blue circles) are indicated with dark grey/light grey lines, respectively. For detailed results, see Figure S 3.

The BayeScan analysis resulted in 391 potential *F*_ST_ outlier loci (SNP_BayeScan). These SNPs, together with those from SNP_adapt, were removed from the SNP_overall to create SNP_neutral (2619 loci). There was almost no overlap between SNP_adapt and SNP_BayeScan (Figure S 2).

### Population genetic diversity

There was a significant interaction effect of habitat type and SNP set on observed heterozygosity (*H*_o_; Table 2). For SNP_neutral, *H*_o_ ranged from 0.21 to 0.30 across all study populations (Table 1). There was no significant difference of *H*_o_ between populations in open and overgrown habitats (Figure 3a; post-hoc test: *p* = 0.91). For SNP_adapt, *H*_o_ ranged from 0.18 to 0.31 across all study populations (Table 1). Importantly, there was a significant difference of *H*_o_ between populations in open and overgrown habitats (post-hoc test: *p* < 0.05), with populations in overgrown habitats exhibiting increased *H*_o_ compared to populations in open habitats (Figure 3a). The random factors region and population accounted for 45.1% and 32.8% of variation in the data for *H*_o_. For the mean of randomised data sets, there was no significant interaction nor habitat effect on *H*_o_ (*p* = 0.58 and *p* = 0.24, respectively) but a significant SNP set effect (*p* < 0.001; Figure S 5a and Figure S 6a). The “permutation *p*-value” for *H*_o_ for the factor habitat type was *p* = 0.05, for SNP set it was *p* = 0.7, and for the interaction it was *p* = 0.04.

**Table 2.**
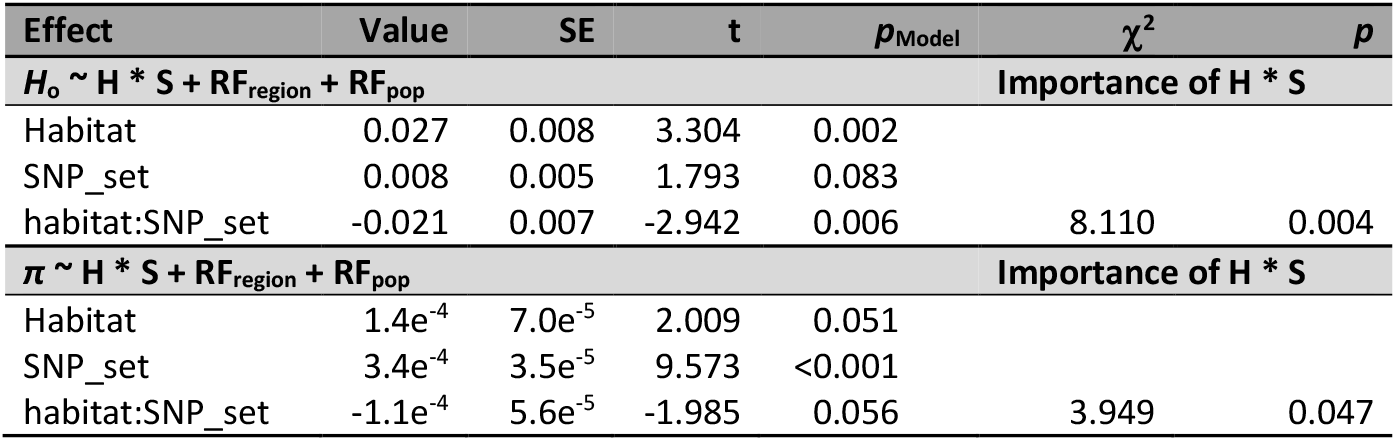
Results of linear mixed effect models for observed heterozygosity (*H*_o_), and nucleotide diversity (*π*) of *Primula veris* populations of the original data set. Fitted parameters (± SE), t-values and significance (*p*_Model_) is given for the full model of each genetic index, *H*_o_ and *π*. Factors: habitat (H): open and overgrown; SNP_set (S): SNP_neutral and SNP_adaptive. RF_region_ and RF_pop_ denote the use of region (Muhu and Saaremaa) and population (1–32) as random factors (RF). The importance of the fixed factor interaction is given as χ^2^ and *p*-values for each genetic measurement. For *H*_o_ and *π*, the importance of the fixed factor interaction is given for the comparison of the full model (with H * S) with the next simplest model (H + S).

**Figure 3.**
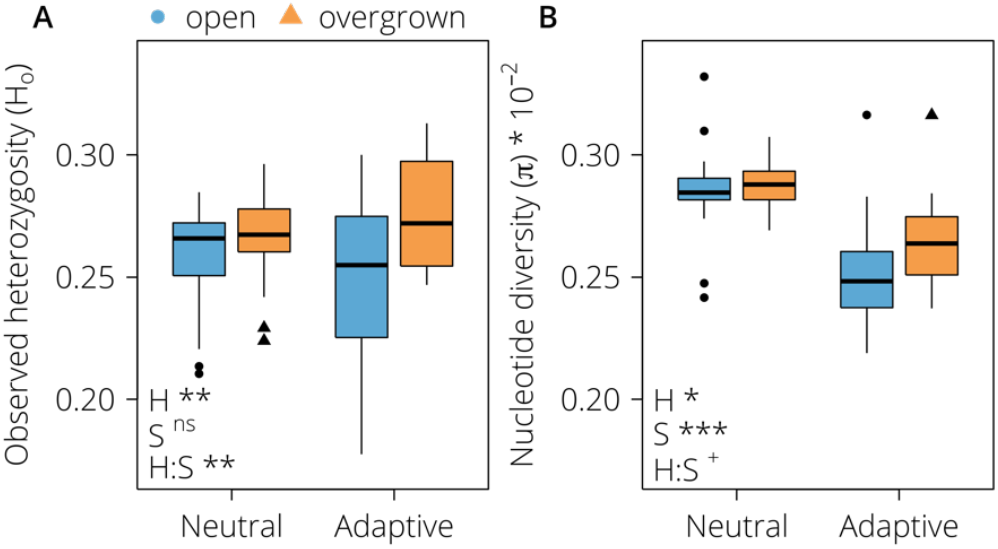
Boxplots of observed heterozygosity (*H*_o_; A) and nucleotide diversity (*π*; B) at putatively neutral and adaptive loci in open (blue, circles) and overgrown (orange, triangles) grasslands. Factors: habitat (H): open and overgrown; SNP_set (S): SNP_neutral and SNP_adaptive. Significance values: ^ns^ *p* > 0.05; ^+^ *p* = 0.056; * *p* ≤ 0.05; ** *p* < 0.01; *** *p* < 0.001.

For nucleotide diversity (*π*), there was a marginally (non-)significant interaction effect of habitat type and SNP set (*p* = 0.056; Table 2). For SNP_neutral, *π* ranged from 0.0024 and 0.0033 across all study populations (Table 1). There was no significant difference of *π* between populations in open and overgrown habitats (Figure 3b; post-hoc test: *p* = 0.97). For SNP_adapt, π ranged from 0.0022 and 0.0032 across all study populations (Table 1). *π* of populations in overgrown habitats was higher than *π* of populations in open habitats, but this difference was not significant (Figure 3b; post-hoc test: *p* = 0.21). The random factors region and population accounted for 0% and 68.4% of variation in the data for *π*. For the mean of randomised data sets, there was no significant interaction nor habitat effect on *π* (*p* = 0.36 and *p* = 0.73, respectively) but a significant SNP set effect (*p* < 0.001; Figure S 5b and Figure S 6b). The “permutation *p*-value” for *π* for the factor habitat type was *p* = 0.14, for SNP set it was *p* = 0, and for the interaction it was *p* = 0.32.

### Population genetic structure and potential gene flow

DAPC of SNP_neutral identified six genetic clusters across the 32 populations with 200 PCs retained (Figure 4a). The first two discriminant functions from DAPC analysis and the hierarchical clustering tree analysis highlighted a differentiation by geographic regions, mainly by the islands Muhu and Saaremaa (Figure 4b,d). Importantly, the separation of genetic clusters was not based on habitat types (Figure 4c,d).

**Figure 4.**
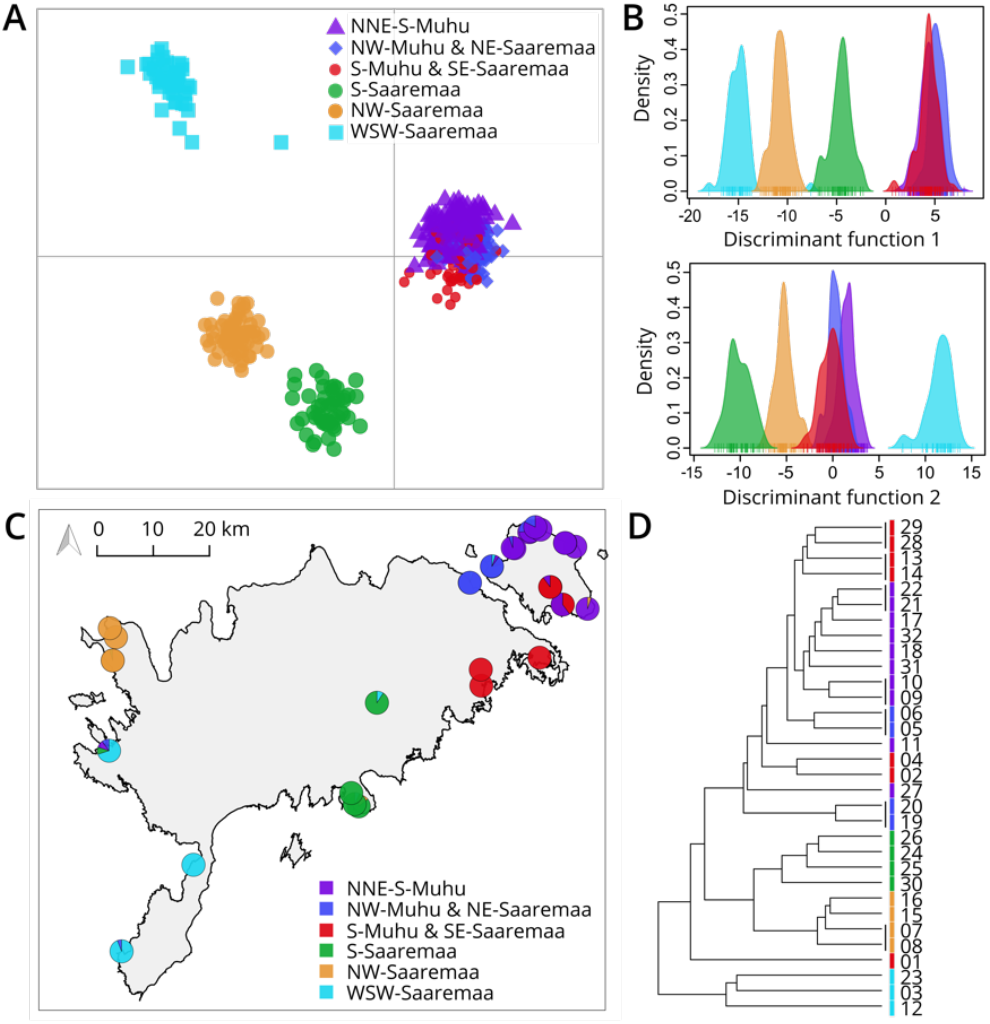
Genetic structure of *Primula veris* populations in the study region. (A) Result of the discriminant analysis of principle components on the putatively neutral SNP set resulting in six geographical clusters (C; named by cardinal directions). The distribution of the six clusters per discriminant function is shown in panel B. (D) Result of the hierarchical clustering tree analysis with population codes (see Table 1) and colour coding by cluster.

Pairwise *F*_ST_ values assessed using SNP_neutral were mostly moderate with an average of 0.10 (± 0.05 SD) ranging from 0.01 to 0.25. We found a positive significant relationship between pairwise *F*_ST_ and geographic distances among all populations (*p*_MCMCglmm_ < 0.001; Table S 2), indicating isolation by distance (IBD; Figure 5). Model fitting showed that values of residual standard error (RSE) of the models roughly reached a plateau between 15 to 30 km in geographic distance between populations (Figure S 7), indicating a potential threshold up to where genetic differentiation is driven by geographic distance and gene flow rather than random (genetic) processes. The habitat type did not explain patterns in genetic differentiation (Table S 2).

**Figure 5.**
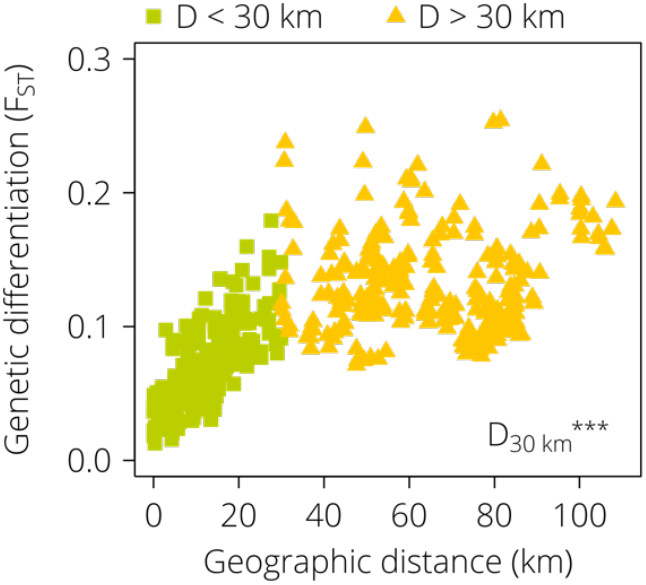
Relationship between genetic differentiation (*F*_ST_) and geographic distance for all possible pairs of 32 *Primula veris* populations, measured at putatively neutral loci. Population pairs with a geographic distance (D) less or equal to 30 km are visualized as green squares. Population pairs with a geographic distance greater than 30 km are given as yellow triangles. The effect of geographic distance on *F*_ST_ is highly significant (*p* < 0.001) up to a threshold of 30 km.

Results of all analyses using SNP_overall were highly similar to when using SNP_neutral. For completeness, these results are presented as supplemental information (Table S 2 – S 4, Figure S 7 – S 10).

## Discussion

Habitat degradation due to abandoned management and related loss and isolation of European semi-natural grasslands (Auffret et al., 2018; Habel et al., 2013) during the last century has been shown to negatively impact the biodiversity of these grasslands, both at the species and the genetic level (e.g. Helm et al., 2006; Picó & Van Groenendael, 2007). Yet, the effect of abandoned management, which can result in gradual overgrowth of grasslands with woody vegetation, on the genetic diversity at adaptive loci of grassland plants has remained unknown. Here, we examined the effect of recent overgrowth of abandoned semi-natural calcareous grasslands (Estonian alvars) on the genetic diversity at putatively neutral and adaptive loci of the perennial herb *Primula veris*. Our study revealed that *P. veris* populations in the new, overgrown habitats likely had a similar level of genetic diversity at putatively neutral loci as in the old, open habitats, despite substantial change in environmental conditions. Genetic diversity at putatively adaptive loci, however, was higher in the new, overgrown compared to the old, open habitats. Neutral genetic structure and gene flow as indicated by neutral genetic differentiation was not (yet) affected by grassland overgrowth. We are among the first to demonstrate how recently changed non-climatic selection pressures are potentially in the process of changing adaptive genetic patterns of wild populations. Most other studies concentrated on the effects of climate on plants (e.g. Dauphin et al., 2020; Sun et al., 2020) or used manipulative experiments in otherwise natural habitats (Laurentino et al., 2020). Hence, our study is an example of potential *in-situ* “adaptation in action”, where genetic diversity at adaptive loci is increasing due to a slow loss of previous genetic adaptations.

Overgrowth of semi-natural grasslands has often a negative impact on genetic diversity of grassland-specialist plants due to a decrease in habitat quality and potential development of barriers for gene flow (Aavik & Helm, 2018; Picó & Van Groenendael, 2007). However, in our study, neutral genetic diversity of *P. veris* was similar in open and overgrown habitats, indicating that habitat and landscape change do not necessarily restrict gene flow among populations of *P. veris*. Still, an environmental impact on the neutral part of a genome might only show after several generations (e.g. Landguth et al., 2010). *Primula veris* is a perennial plant with an average lifespan of up to 50 years and can persist with reduced reproduction even when the environment has changed due to overgrowth (Ehrlén & Lehtilä, 2002). Hence, genetic diversity of such populations potentially reflects the state before their habitats started to change (Reinula, 2018). In addition, heterozygosity indices, as used in our study, have been suggested to respond more slowly to environmental change as compared to, for example, measures of inbreeding (Lloyd, Campbell, & Neel, 2013; Lowe, Boshier, Ward, Bacles, & Navarro, 2005). Deschepper et al. (2017), who examined patterns of neutral genetic diversity of *P. veris* in grassland and forest populations in Belgium, also reported no significant difference in *H*_*o*_ between habitat types. Yet, their study system, i.e. forests, represents a late-successional stage, whereas ours represents a mid-successional stage (shrubby overgrowth). Consequently, neutral genetic patterns of *P. veris* might need a very long time (up to centuries) to reach an equilibrium with the new overgrown environment.

In contrast to genetic diversity at putatively neutral loci, genetic diversity assessed at putatively adaptive loci differed between habitat types in our study. Populations in recently overgrown grasslands showed higher genetic diversity at putatively adaptive loci compared to populations in open grasslands (Figure 3a,b). This is most likely caused by changed selection pressures in the new, overgrown habitats compared to the old, open habitats. The open grasslands used in our study have been in an open state for several hundreds of years, whereas ongoing overgrowth started only about 90 years ago (Helm et al., 2006). Hence, populations of *P. veris* in open grasslands have experienced homogenous selection pressures for a very long time, which increased the frequencies of alleles beneficial for an open habitat and led to reduced heterozygosity and genetic diversity at adaptive loci. The majority of putatively adaptive SNPs (53 out of 77) exhibited a smaller average beneficial (i.e. major) AF for open habitat conditions in populations from the new, overgrown compared to those from the old habitat. This led to an increase in heterozygosity and genetic diversity in populations of the new, overgrown habitats (Figure 2). Considering *P. veris’* potential longevity, this indicates that populations in overgrown habitats are still adapting to their new selection pressures (e.g. lower light availability, reduced or altered pollinator community), i.e. many alleles potentially beneficial for the new, overgrown habitat are still far from fixation (i.e. homozygosity). Consequently, the genetic diversity at putatively adaptive loci of *P. veris* populations in overgrown grasslands has not yet been reduced. A similar pattern was found in the conifer tree *Pinus cembra*, where populations in the core of the current niche exhibited a decreased genetic diversity at adaptive loci, but no difference in the one at neutral loci, compared to populations at the niche margin, which most likely present unstable or novel habitats (Dauphin et al., 2020).

An increase of non-beneficial alleles for the open habitat in overgrown populations, and thus an increase in heterozygosity at putatively adaptive loci in overgrown populations, can be achieved in two ways: (1) the alternative alleles at SNP loci were either already present in lower frequencies in populations of open habitats (i.e. standing genetic variation), or (2) arrived to overgrowing grasslands by gene flow, before they were subject to positive selection in the overgrown habitat. As shown in Figure S 3, beneficial alleles in populations of the open habitat were rarely fixed, strongly pointing towards the importance of standing genetic variation. This implies that even populations with reduced genetic diversity at putatively adaptive loci (e.g. in open grasslands) may possess the ability to react to habitat changes due to the low, but crucial amount of standing genetic variation (e.g. Morris, Bowles, Allen, Jamniczky, & Rogers, 2018). Still, we found gene flow as indicated by genetic differentiation potentially spreading alleles between habitats which, in theory, can contribute to the increased genetic diversity at adaptive loci in overgrown populations. The potential gene flow distances found in our study (up to 30 km; Figure 4, Figure 5) stand in marked contrast to the potential dispersal ranges of pollen and seeds of *P. veris* for which very short distances up to 12 m and 0.5 m, respectively, have been reported (Antrobus & Lack, 1993; Richards & Ibrahim, 1978). Yet, pollinating insects of *P. veris* have been shown to occasionally travel up to 2 km (Kreyer, Oed, Walther-Hellwig, & Frankl, 2004; Zurbuchen, Bachofen, Müller, Hein, & Dorn, 2010) and dispersal distances are often underestimated in ecological studies (e.g. Bullock, Shea, & Skarpaas, 2006). Historical rotational grazing of domestic animals and movement of wild animals (e.g. deer, moose, wild boar) might also have facilitated seed dispersal for longer distances in *P. veris* which is not adapted to zoochory (Plue, Aavik, & Cousins, 2019). Overall, standing genetic variation but also gene flow are capable of supplying *P. veris* populations undergoing habitat changes with new or alternative alleles fostering adaptation to new habitat conditions in our study region.

Importantly, the fact that we detected a significant effect of habitat type on genetic diversity (*H*_*o*_) when using the putatively adaptive SNP set but not with the neutral SNP set suggests the need to examine genetic diversity at neutral and adaptive loci separately when studying the genetic response of plant species to environmental changes. Besides, in the overall SNP set (Figure S 10), there was no significant effect of habitat type on genetic diversity, which indicates that genetic diversity of an overall SNP set represents rather neutral genetic patterns (Dauphin et al., 2020). In conservation genetics and restoration, most assessments have been based on overall or neutral genetic diversity so far (González et al., 2019; Wei & Jiang, 2020), even though its stand-alone importance is questionable (Teixeira & Huber, 2021) and it is likely the putatively adaptive regions of a genome that are important for the fate of a population in a changed environment.

The different effects of habitat type on genetic diversity at putatively neutral and adaptive loci could partly be due to the unevenness in the number of loci at neutral and putatively adaptive regions (2,619 versus 77, respectively). However, the ddRADseq procedure used in our study should result in a “random” and representative subset of both, neutral and adaptive SNPs. Here, we were not particularly interested in the actual molecular mechanisms underlying adaptation, but in general patterns at putatively adaptive loci, which should also become visible with 77 SNPs if these patterns are of substantial nature. Additionally, the reliability of the detected SNPs can be assumed high, because the genotyping error for our SNPs was low (0.004) and the SNP sets were identified using a draft genome (Hoban et al., 2016), which covers about 63% of the whole 479.22 Mb genome of the study species *P. veris* (Nowak et al., 2015).

When using linear and categorical EAA to test effects of habitat type on genetic diversity, we must consider a potential circularity issue. Nevertheless, the results of the randomised data sets indicate that, for *H*_*o*_, the habitat type related results (including the interaction with SNP set) of the original data are likely not concerned by circularity. Yet, this is the case for *π*. Potentially, since our study sites are just in the beginning of a habitat change process (i.e. in terms of perennial plants like *P. veris*), more time might be needed until a stronger habitat change effect can be detected by *π*. Importantly, even though the number of putatively adaptive loci using randomised data sets is in the range of putatively adaptive loci found using the original data set, the putatively adaptive loci from the original data set do not show a random pattern with respect to their change in beneficial AF (Figure S 4). In addition, applying the same mixed-effect model statistic to the mean randomised data set, for both *H*_*o*_ and *π*, there is no habitat type or interaction effect anymore (Figure S 5), as can be expected when breaking the apparent link between genotype and habitat type sample assignment of the original data set. Therefore, caution is needed when interpreting the results, especially those using *π*, but the overall conclusion of potential early stages of “adaption in action” processes remains. To our knowledge, there is no previous study applying EAA to examine putatively neutral and adaptive genetic diversity that used permutations for validation. In contrast, such validations are common practice in genome wide association studies (e.g. Kreiner, Tranel, Weigel, Stinchcombe, & Wright, 2021; Tibbs Cortes, Zhang, & Yu, 2021) which, together with our results, might call for their wider application in landscape genomic analyses.

In addition to genetic diversity, population size is another factor that is affected by habitat change, and which also directly influences genetic diversity (Leimu et al., 2006). The census sizes of our *P. veris* populations were similar in open and overgrown habitats and were not associated with genetic diversity at neutral and adaptive loci. In contrast to our results, a meta-analysis showed a clear relationship between population size and (neutral) genetic diversity, including long-lived and self-incompatible plant species, such as *P. veris* (Leimu et al., 2006). Populations in our study exhibited a minimum of about 100 individuals, which is larger than the overall census for small populations in the study by Leimu et al. (2006). Consequently, our *P. veris* populations might be still sufficiently large to counteract population size driven effects on genetic diversity.

## Conclusions

Our landscape genomic investigation of *Primula veris* in Estonian semi-natural grasslands is one of the first to demonstrate the potential effects of land use change on the genetic diversity at putatively adaptive loci of *in-situ* wild plant species. We show that the effect of recent overgrowth of grasslands might not be genetically manifested when considering neutral SNPs, even after almost a century of ongoing environmental changes. Yet, genetic diversity assessed at putatively adaptive loci was higher in populations in overgrown compared to open habitats, most probably due to allele frequency changes of standing genetic variation. Thus, even populations in degraded and fragmented habitats may possess the ability to adapt to habitat changes due to their important standing genetic variation in addition to potential allele immigration due to gene flow. However, permutation analyses indicated that caution is needed when interpreting results based on linear and categorical EAA, especially when working with nucleotide diversity.

For perennial long-lived plant species, such as *P. veris*, long time spans might pass before habitat change can be detected at neutral regions of the genome whereas habitat effects at adaptive loci could be noticeable faster. Consequently, a repeated monitoring of genetic diversity at both neutral and adaptive loci and further investigations of contemporary gene flow at different spatial scales would be highly valuable to identify potential genetic consequences of recent and ongoing environmental change in natural and semi-natural habitats. In addition, extending our results to whole-genome and targeted sequencing approaches would be vital to reliably identify genes and gene networks putatively involved in the adaptation of *P. veris* to habitat change, and to assess the relative importance of loss of previous and gain of new genetic adaptations to the altered environment.

## Acknowledgements

We thank the Genetic Diversity Centre Zurich (GDC) for laboratory support, the Functional Genomic Centre Zurich (FGCZ) for Illumina sequencing, A. Rogivue for introducing the lead author to EAA, and B. Dauphin for extracting climate data and support in EAAs. We are grateful for financial support from the Estonian Research Council (MOBJD427, PUT589 and PRG874), the COST “G-BIKE” action (CA18134), the European Regional Development Fund (Centre of Excellence EcolChange), and European Commission LIFE+ Nature program (LIFE13NAT/EE/000082). We also thank three anonymous reviewers for their valuable comments and suggestions on previous versions of this manuscript.

## Data Accessibility

Sequence data used in this study will be made available at the European Nucleotide Archive (ENA) upon acceptance (ERS5253979 – ERS5254546). R-scripts, genotypic and environmental data will be provided at the Dryad Digital Repository upon acceptance (xxx).

## Author Contributions

T.A., S.T. and A.H designed the conceptual approach and carried out field work. S.T. and I.R. conducted laboratory work. S.T. and N.Z. performed bioinformatic analyses. S.T. and C.R. analysed the data. R.H. contributed with discussing the results in a broader ecological context. S.T. wrote the manuscript with major contributions from C.R. All authors read, commented and approved the final version of the manuscript.

## Supplemental Information

### Supplemental Methods

Figure S 1 Environmental variables and their response to habitat type (open – overgrown).

Figure S 2 Venn diagram of shared putatively adaptive loci by different methods.

Figure S 3 SNP allele frequencies and their behaviour in open and overgrown habitats.

Figure S 4 Proportion of putatively adaptive SNPs with de- and increasing beneficial AF of the original and randomised data sets.

Figure S 5 Mean of genetic and nucleotide diversity (*H*_o_ and *π*) at putatively neutral and adaptive SNPs of 100 randomised data sets.

Figure S 6 Genetic and nucleotide diversity (*H*_o_ and *π*) at putatively neutral and adaptive SNPs per population of 100 randomised data sets.

Figure S 7 Residual standard error results for IBD analyses, measured using the overall (3,084 loci) and neutral (2,619 loci) SNP sets.

Figure S 8 Genetic structure of *Primula veris* populations in the study area measured using the overall set of loci (3,084 loci).

Figure S 9 Isolation by distance pattern measured at the overall (3,084 SNPs) set of loci for *Primula veris* populations.

Figure S 10 Genetic and nucleotide diversity (*H*_o_ and *π*) at the overall set of SNPs (3,084 loci).

Table S 1 Number of associations between SNPs and environmental variables.

Table S 2 Results of generalized mixed effect models for the effect of geographic distance and habitat type distance on pairwise genetic differentiation (*F*_ST_) for the neutral (2,619 loci) and overall (3,084 loci) SNP sets.

Table S 3 Genetic diversity measures using the overall set of SNPs (3,084 loci) for the studied populations of *Primula veris*.

Table S 4 Results of linear mixed effect models for *H*_o_, *π* for the overall SNP set (3,084 loci) of the studied *Primula veris* populations.

## References

Aavik, T., & Helm, A. (2018). Restoration of plant species and genetic diversity depends on landscape-scale dispersal. Restoration Ecology. 26, S92–S102. doi: 10.1111/rec.12634

Aavik, T., Thetloff, M., Träger, S., Hernández-Agramonte, I. M., Reinula, I., & Pärtel, M. (2019). Delayed and immediate effects of habitat loss on the genetic diversity of the grassland plant Trifolium montanum. Biodiversity and Conservation, 28, 3299–3319. doi: 10.1007/s10531-019-01822-8

Antrobus, S., & Lack, A. J. (1993). Genetics of colonizing and established populations of Primula veris. Heredity, 71, 252–258. doi: 10.1038/hdy.1993.133

Auffret, A. G., Kimberley, A., Plue, J., & Waldén, E. (2018) Super-regional land-use change and effects on the grassland specialist flora. Nature communications, 9, 3464. doi: 10.1038/s41467-018-05991-y.

Barrett, R. D. H., & Schluter, D. (2008). Adaptation from standing genetic variation. Trends in Ecology and Evolution, 23, 38–44. doi: 10.1016/j.tree.2007.09.008

Benjamini, Y., & Hochberg, Y. (1995). Controlling the false discovery rate: a practical and powerful approach to multiple testing. Journal of the Royal Statistical Society: Series B (Methodological), 57, 289–300. doi: 10.1111/j.2517-6161.1995.tb02031.x

Bilska, K., & Szczecińska, M. (2016). Comparison of the effectiveness of ISJ and SSR markers and detection of outlier loci in conservation genetics of Pulsatilla patens populations. PeerJ, 4, e2504. doi: 10.7717/peerj.2504

Bolger, A. M., Lohse, M., & Usadel, B. (2014). Trimmomatic: a flexible trimmer for Illumina sequence data. Bioinformatics, 30, 2114–2120. doi: 10.1093/bioinformatics/btu170

Brys, R., & Jacquemyn, H. (2009). Biological Flora of the British Isles: Primula veris L. Journal of Ecology, 97, 581–600. doi: 10.1111/j.1365-2745.2009.01495.x

Bullock, J. M., Shea, K., & Skarpaas, O. (2006). Measuring plant dispersal: an introduction to field methods and experimental design. Plant Ecology, 186, 217–234. doi: 10.1007/s11258-006-9124-5

Catchen, J. M., Amores, A., Hohenlohe, P., Cresko, W., Postlethwait, J. H., & De Koning, D.-J. (2011). Stacks: building and genotyping loci de novo from short-read sequences. G3: Genes|Genomes||Genetics, 1(3), 171–182. doi: 10.1534/g3.111.000240

Catchen, J. M., Hohenlohe, P. A., Bassham, S., Amores, A., & Cresko, W. A. (2013). Stacks: an analysis tool set for population genomics. Molecular Ecology, 22, 3124–3140. doi: 10.1111/mec.12354.Stacks

Caye, K., Jumentier, B., Lepeule, J., & François, O. (2019). LFMM 2: Fast and accurate inference of gene-environment associations in genome-wide studies. Molecular Biology and Evolution, 36, 852–860. doi: 10.1093/molbev/msz008

Chambers, J. M. (1992). Linear models. In J. M. Chambers & T. J. Hastie (Eds.), Statistical Models in S. Wadsworth & Brooks/Cole.

Cheptou, P.-O., Hargreaves, A. L., Bonte, D., & Jacquemyn, H. (2017). Adaptation to fragmentation: evolutionary dynamics driven by human influences. Philosophical Transactions of the Royal Society B: Biological Sciences, 372, 20160037. doi: 10.1098/rstb.2016.0037

Coop, G., Witonsky, D., Di Rienzo, A., & Pritchard, J. K. (2010). Using environmental correlations to identify loci underlying local adaptation. Genetics, 185, 1411–1423. doi: 10.1534/genetics.110.114819

Cousins, S. A. O., Auffret, A. G., Lindgren, J., & Tränk, L. (2015). Regional-scale land-cover change during the 20th century and its consequences for biodiversity. AMBIO, 44, 17– 27. doi: 10.1007/s13280-014-0585-9

Danecek, P., Auton, A., Abecasis, G., Albers, C. A., Banks, E., DePristo, M. A., … Durbin, R. (2011). The variant call format and VCFtools. Bioinformatics, 27, 2156–2158. doi: 10.1093/bioinformatics/btr330

Dauphin, B., Wüest, R. O., Brodbeck, S., Zoller, S., Fischer, M. C., Holderegger, R., … Rellstab, C. (2020). Disentangling the effects of geographic peripherality and habitat suitability on neutral and adaptive genetic variation in Swiss stone pine. Molecular Ecology, 29, 1972–1989. doi: 10.1111/mec.15467

Davey, J. W., Hohenlohe, P. A., Etter, P. D., Boone, J. Q., Catchen, J. M., & Blaxter, M. L. (2011). Genome-wide genetic marker discovery and genotyping using next-generation sequencing. Nature Reviews Genetics, 12, 499–510. doi: 10.1038/nrg3012

de Mita, S., Thuillet, A.-C., Gay, L., Ahmadi, N., Manel, S., Ronfort, J., & Vigouroux, Y. (2013). Detecting selection along environmental gradients: analysis of eight methods and their effectiveness for outbreeding and selfing populations. Molecular Ecology, 22, 1383– 1399. doi: 10.1111/mec.12182

Deschepper, P., Brys, R., & Jacquemyn, H. (2018). The impact of flower morphology and pollinator community composition on pollen transfer in the distylous Primula veris. Botanical Journal of the Linnean Society, 186, 414–424. doi: 10.1093/botlinnean/box097

DiLeo, M. F., Holderegger, R., & Wagner, H. H. (2018). Contemporary pollen flow as a multiscale process: Evidence from the insect-pollinated herb Pulsatilla vulgaris. Journal of Ecology, 0–3. doi: 10.1111/1365-2745.12992

Ehrlén, J., & Lehtilä, K. (2002). How perennial are perennial plants? Oikos, 98(2), 308–322.

Exposito-Alonso, M., 500 Genomes Field Experiment Team, Burbano, H. A., Bossdorf, O., Nielsen, R., & Weigel, D. (2019). Natural selection on the Arabidopsis thaliana genome in present and future climates. Nature, 573, 126–129. doi: 10.1038/s41586-019-1520-9

EWS. (2020). Estonian Weather Service. http://www.ilmateenistus.ee

Foll, M., & Gaggiotti, O. (2008). A genome-scan method to identify selected loci appropriate for both dominant and codominant markers: a Bayesian perspective. Genetics, 180, 977–993. doi: 10.1534/genetics.108.092221

François, O., Martins, H., Caye, K., & Schoville, S. D. (2016). Controlling false discoveries in genome scans for selection. Molecular Ecology, 25, 454–469. doi: 10.1111/mec.13513

Frankham, R. (2005). Genetics and extinction. Biological Conservation, 126, 131–140. doi: 10.1016/j.biocon.2005.05.002

Garrison, E., & Marth, G. (2012). Haplotype-based variant detection from short-read sequencing. ArXiv, 1207.3907. doi: 1207.3907 [q-bio.GN]

González, A. V., Gómez-Silva, V., Ramírez, M. J., Fontúrbel, F. E. (2019). Meta-analysis of the differential effects of habitat fragmentation and degradation on plant genetic diversity. Conservation Biology, 34, 711–720. doi: 10.1111/cobi.13422

Habel, J. C., Dengler, J., Janišová, M., Török, P., Wellstein, C., & Wiezik, M. (2013). European grassland ecosystems: threatened hotspots of biodiversity. Biodiversity and Conservation, 22, 2131–2138. doi: 10.1007/s10531-013-0537-x

Hadfield, J. D. (2010). MCMC methods for multi-response generalized linear mixed models: the MCMCglmm R package. Journal of Statistical Software, 33, 1–22. doi: 10.1002/ana.22635

Hamrick, J. L., & Godt, M. J. W. (1996). Effects of life history traits on genetic diversity in plant species. Philosophical Transactions: Biological Sciences, 351, 1291–1298. doi: 10.1098/rstb.1996.0112

Helm, A. (2019). Large-scale restoration of Estonian alvar grasslands: impact on biodiversity and ecosystem services - Final report of the Action D.1. Biodiversity monitoring for project LIFE to Alvars (LIFE13NAT/EE/000082). https://life.envir.ee/sites/default/files/pictures/LIFE_to_Alvars_Report_Biodiversity_monitoring_submitted.pdf

Helm, A., Hanski, I., & Pärtel, M. (2006). Slow response of plant species richness to habitat loss and fragmentation. Ecology Letters, 9, 72–77. doi: 10.1111/j.1461-0248.2005.00841.x

Hoban, S., Kelley, J. L., Lotterhos, K. E., Antolin, M. F., Bradburd, G., Lowry, D. B., … Whitlock, M. C. (2016). Finding the genomic basis of local adaptation: pitfalls, practical solutions, and future directions. The American Naturalist, 188, 379–397. doi: 10.1086/688018

Honnay, O., & Jacquemyn, H. (2007). Susceptibility of common and rare plant species to the genetic consequences of habitat fragmentation. Conservation Biology, 21, 823–831. doi: 10.1111/j.1523-1739.2006.00646.x

Hooftman, D. A. P., & Bullock, J. M. (2012). Mapping to inform conservation: a case study of changes in semi-natural habitats and their connectivity over 70 years. Biological Conservation, 145, 30–38. doi: 10.1016/j.biocon.2011.09.015

IPBES. (2018). Summary for policymakers of the regional assessment report on biodiversity and 24 ecosystem services for Europe and Central Asia of the Intergovernmental Science-Policy Platform on 25 Biodiversity and Ecosystem Services. Fischer, M., M. Rounsevell, A. Torre-Marin Rando, A. Mader, A. Church, M. Elbakidze, V. Elias, H. T. P. A. Harrison, J. Hauck, B. Martín-López, I. Ring, C. Sandström, I. Sousa Pinto, P. Visconti, and N. E. Zimmermann: IPBES secretariat, Bonn, Germany.

IPCC. (2019). Climate Change and Land: an IPCC special report on climate change, desertification, land degradation, sustainable land management, food security, and greenhouse gas fluxes in terrestrial ecosystems. P. R. Shukla, J. Skea, E. Calvo Buendia, V. Masson-Delmotte, H.-O. Pörtner, D. C. Roberts, P. Zhai, R. Slade, S. Connors, R. van Diemen, M. Ferrat, E. Haughey, S. Luz, S. Neogi, M. Pathak, J. Petzold, J. Portugal Pereira, P. Vyas, E. Huntley, K. Kissick, M. Belkacemi, J. Malley, (eds.).

Jombart, T. (2008). Adegenet: a R package for the multivariate analysis of genetic markers. Bioinformatics, 24, 1403–1405. doi: 10.1093/bioinformatics/btn129

Jombart, T., Devillard, S., & Balloux, F. (2010). Discriminant analysis of principal components: a new method for the analysis of genetically structured populations. BMC Genetics, 11. doi: https://doi.org/10.1186/1471-2156-11-94

Kamvar, Z. N., Tabima, J. F., Everhart, S. E., Brooks, J. C., Krueger-Hadfield, S. A., Sotka, E., … Grünwald, N. J. (2019). poppr - Genetic analysis of populations with mixed reproduction.

Karger, D. N., Conrad, O., Böhner, J., Kawohl, T., Kreft, H., Soria-Auza, R. W., … Kessler, M. (2017). Climatologies at high resolution for the earth’s land surface areas. Scientific Data, 4, 170122. doi: 10.1038/sdata.2017.122

Kreiner, J. M., Tranel, P. J., Weigel, D., Stinchcombe, J. R., & Wright, S. I. (2021). The genetic architecture and population genomic signatures of glyphosate resistance in Amaranthus tuberculatus. Molecular Ecology. doi: 10.1111/mec.15920

Kreyer, D., Oed, A., Walther-Hellwig, K., & Frankl, R. (2004). Are forests potential landscape barriers for foraging bumblebees? Landscape scale experiments with Bombus terrestris agg. and Bombus pascuorum (Hymenoptera, Apidae). Biological Conservation, 116, 111–118. doi: 10.1016/S0006-3207(03)00182-4

Kuznetsova, A., Brockhoff, P. B., & Christensen, R. H. B. (2017). lmerTest Package: Tests in linear mixed effects models. Journal of Statistical Software, 82, 1–26. doi: 10.18637/jss.v082.i13

Landguth, E. L., Cushman, S. A., Schwartz, M. K., McKelvey, K. S., Murphy, M., & Luikart, G. (2010). Quantifying the lag time to detect barriers in landscape genetics. Molecular Ecology, 19, 4179–4191. doi: 10.1111/j.1365-294X.2010.04808.x

Laurentino, T. G., Moser, D., Roesti, M., Ammann, M., Frey, A., Ronco, F., Kueng, B., & Berner, D. (2020). Genomic release-recapture experiment in the wild reveals within-generation polygenic selection in stickleback fish. Nature Communications, 11, 1928. doi: 10.1038/s41467-020-15657-3

Lehtilä, K., Dahlgren, J. P., Garcia, M. B., Leimu, R., Syrjänen, K., & Ehrlén, J. (2016). Forest succession and population viability of grassland plants: long repayment of extinction debt in Primula veris. Oecologia, 181, 125–135. doi: 10.1007/s00442-016-3569-6

Leimu, R., Mutikainen, P., Koricheva, J., & Fischer, M. (2006). How general are positive relationships between plant population size, fitness and genetic variation? Journal of Ecology, 94, 942–952. doi: 10.1111/j.1365-2745.2006.01150.x

Lenth, R. V. (2016). Least-squares means: The R package lsmeans. Journal of Statistical Software, 69, 1–33. doi: 10.18637/jss.v069.i01

Li, H. (2013). Aligning sequence reads, clone sequences and assembly contigs with BWA-MEM. ArXiv, 1303.3997. doi: 1303.3997 [q-bio.GN]

Lloyd, M. W., Campbell, L., & Neel, M. C. (2013). The power to detect recent fragmentation events using genetic differentiation methods. PLoS ONE, 8, e63981. doi: 10.1371/journal.pone.0063981

Lotterhos, K. E., & Whitlock, M. C. (2015). The relative power of genome scans to detect local adaptation depends on sampling design and statistical method. Molecular Ecology, 24, 1031–1046. doi: 10.1111/mec.13100

Lowe, A. J., Boshier, D., Ward, M., Bacles, C. F. E., & Navarro, C. (2005). Genetic resource impacts of habitat loss and degradation; reconciling empirical evidence and predicted theory for neotropical trees. Heredity, 95, 255–273. doi: 10.1038/sj.hdy.6800725

Metzker, M. L. (2010). Sequencing technologies — the next generation. Nature Reviews Genetics, 11, 31–46. doi: 10.1038/nrg2626

Milot, E., Béchet, A., & Maris, V. (2020) The dimensions of evolutionary potential in biological conservation. Evolutionary Applications, 13, 1363–1379. doi: 10.1111/eva.12995

Morris, M. R. J., Bowles, E., Allen, B. E., Jamniczky, H. A., & Rogers, S. M. (2018). Contemporary ancestor? Adaptive divergence from standing genetic variation in Pacific marine threespine stickleback. BMC Evolutionary Biology, 18, 113. doi: 10.1186/s12862-018-1228-8

Nei, M., Suzuki, Y., & Nozawa, M. (2010). The neutral theory of molecular evolution in the genomic era. Annual Review of Genomics and Human Genetics, 11, 265–289. doi: 10.1146/annurev-genom-082908-150129

Nowak, M. D., Russo, G., Schlapbach, R., Huu, C., Lenhard, M., & Conti, E. (2015). The draft genome of Primula veris yields insights into the molecular basis of heterostyly. Genome Biology, 16, 12. doi: 10.1186/s13059-014-0567-z

Peakall, R., & Smouse, P. E. (2012). GenAlEx 6.5: genetic analysis in Excel. Population genetic software for teaching and research — an update. Bioinformatics, 28, 2537–2539. doi: 10.1093/bioinformatics/bts460

Peterson, B. K., Weber, J. N., Kay, E. H., Fisher, H. S., & Hoekstra, H. E. (2012). Double digest RADseq: an inexpensive method for de novo SNP discovery and genotyping in model and non-model species. PLoS ONE, 7. doi: 10.1371/journal.pone.0037135

Picó, F. X., & Van Groenendael, J. (2007). Large-scale plant conservation in European semi-natural grasslands: a population genetic perspective. Diversity and Distributions, 13, 920–926. doi: 10.1111/j.1472-4642.2007.00349.x

Plue, J., Aavik, T., & Cousins, S. A. O. (2019). Grazing networks promote plant functional connectivity among isolated grassland communities. Diversity and Distributions, 25, 102–115. doi: 10.1111/ddi.12842

Pollard, E. (1977). A method for assessing changes in the abundance of butterflies. Biological Conservation, 12, 115–134.

Prangel, E. (2017). The provisioning of ecosystem services on open and successional alvar grasslands (Msc thesis). University of Tartu, Tartu.

Puritz, J. B., Hollenbeck, C. M., & Gold, J. R. (2014). dDocent: a RADseq, variant-calling pipeline designed for population genomics of non-model organisms. PeerJ, 2, e431. doi: 10.7717/peerj.431

Puritz, J. B., Matz, M. V., Toonen, R. J., Weber, J. N., Bolnick, D. I., & Bird, C. E. (2014). Demystifying the RAD fad. Molecular Ecology, 23, 5937–5942. doi: 10.1111/mec.12965

R Development Core Team. (2017). R: A language and environment for statistical computing (3.2.3). Vienna: R Foundation for Statistical Computing.

Radwan, J., & Babik, W. (2012). The genomics of adaptation. Proceedings of the Royal Society B: Biological Sciences, 279, 5024–5028. doi: 10.1098/rspb.2012.2322

Reinula, I. (2018). The impact of landscape change on the genetic diversity of the grassland plant Primula veris (Msc thesis). University of Tartu, Tartu.

Rellstab, C., Gugerli, F., Eckert, A. J., Hancock, A. M., & Holderegger, R. (2015). A practical guide to environmental association analysis in landscape genomics. Molecular Ecology, 24, 4348–4370. doi: 10.1111/mec.13322

Rellstab, C., Zoller, S., Walthert, L., Lesur, I., Pluess, A. R., Graf, R., … Gugerli, F. (2016). Signatures of local adaptation in candidate genes of oaks (Quercus spp.) with respect to present and future climatic conditions. Molecular Ecology, 25, 5907–5924. doi: 10.1111/mec.13889

Richards, A. J., & Ibrahim, H. (1978). Estimation of neighbourhood size in two populations of Primula veris. In A. J. Richards (Ed.), The pollination of flowers by insects (pp. 165–174). London, UK: Academic Press.

Rosenberg, N. A., & Jakobsson, M. (2008). The relationship between homozygosity and the frequency of the most frequent allele. Genetics, 179, 2027–2036. doi: 10.1534/genetics.107.084772

Rousset, F., Lopez, J., & Belkhir, K. (2017). genepop - Population genetic data analysis using genepop.

Slatkin, M. (1985). Gene flow in natural populations. Annual Review of Ecology and Systematics, 16, 393–430. doi: doi:10.1146/annurev.es.16.110185.002141

Smith, A. L., Hodkinson, T. R., Villellas, J., Catford, J. A., Csergő, A. M., Blomberg, S. P., … Buckley, Y. M. (2020). Global gene flow releases invasive plants from environmental constraints on genetic diversity. Proceedings of the National Academy of Sciences, 117, 4218–4227. doi: 10.1073/pnas.1915848117

Stoeckel, S., Grange, J., Fernández-Manjarres, J. F., Bilger, I., Frascaria-Lacoste, N., & Mariette, S. (2006). Heterozygote excess in a self-incompatible and partially clonal forest tree species — Prunus avium L. Molecular Ecology, 15, 2109–2118. doi: 10.1111/j.1365-294X.2006.02926.x

Sun, Y., Bossdorf, O., Grados, R. D., Liao, Z., & Müller-Schärer, H. (2020). Rapid genomic and phenotypic change in response to climate warming in a widespread plant invader. Global Change Biology, 26, 6511–6522. doi: 10.1111/gcb.15291

Teixeira, J. C., & Huber, C. D. (2021). The inflated significance of neutral genetic diversity in conservation genetics. Proceedings of the National Academy of Sciences, 118. doi: 10.1073/pnas.2015096118

Tibbs Cortes, L., Zhang, Z., & Yu, J. (2021). Status and prospects of genome-wide association studies in plants. The Plant Genome, 14. doi: 10.1002/tpg2.20077

Tewksbury, J. J., Levey, D. J., Haddad, N. M., Sargent, S., Orrock, J. L., Weldon, A., … Townsend, P. (2002). Corridors affect plants, animals, and their interactions in fragmented landscapes. Proceedings of the National Academy of Sciences, 99, 12923–12926. doi: 10.1073/pnas.202242699

Van Strien, M. J., Holderegger, R., & Van Heck, H. J. (2015). Isolation-by-distance in landscapes: considerations for landscape genetics. Heredity, 114, 27–37. doi: 10.1038/hdy.2014.62

Wedderburn, F., & Richards, A. J. (1990). Variation in within-morph incompatibility inhibition sites in heteromorphic Primula L. New Phytologist, 116, 149–162.

Wegmann Lab. (2019). TIGER - tools to estimate genotyping errors (Version 1.0). https://bitbucket.org/wegmannlab/tiger/wiki/Home

Wei, X., & Jiang, M. (2020). Meta-analysis of genetic representativeness of plant populations under ex situ conservation in contrast to wild source populations. Conservation Biology. doi: 10.1111/cobi.13617

Westergaard, K. B., Zemp, N., Bruederle, L. P., Stenøien, H. K., Widmer, A., & Fior, S. (2019). Population genomic evidence for plant glacial survival in Scandinavia. Molecular Ecology, 28, 818–832. doi: 10.1111/mec.14994

Wilson, J. B., Peet, R. K., Dengler, J., & Pärtel, M. (2012). Plant species richness: the world records. Journal of Vegetation Science, 23, 796–802. doi: 10.1111/j.1654-1103.2012.01400.x

Xuereb, A., Kimber, C. M., Curtis, J. M. R., Bernatchez, L., & Fortin, M. J. (2018). Putatively adaptive genetic variation in the giant California sea cucumber (Parastichopus californicus) as revealed by environmental association analysis of restriction-site associated DNA sequencing data. Molecular Ecology, 27, 5035–5048. doi: 10.1111/mec.14942

Zurbuchen, A., Bachofen, C., Müller, A., Hein, S., & Dorn, S. (2010). Are landscape structures insurmountable barriers for foraging bees? A mark-recapture study with two solitary pollen specialist species. Apidologie, 41, 497–508. doi: 10.1051/apido/2009084

