## Supporting Information for "The effect of recent habitat change on genetic diversity at putatively adaptive and neutral loci in *Primula veris* in semi-natural grasslands"

### Supplemental Methods

#### Adapter annealing

In our study, we used two different restriction enzymes, EcoRI and Taq$^{\alpha}$I, since they resulted in the highest expected number of restriction site associated fragments according to SimRAD simulations (Lepais & Weir, 2014). In this simulation, different combinations of restriction enzymes with EcoRI were tested (e.g. EcoRI – TaqI, EcoRI – MseI, EcoRI – SbfI, EcoRI – SphI, EcoRI – MspI, EcoRI – PseI).

EcoRI barcode adapters for individual barcoding were used according to Peterson et al. (2012). The 48 EcoRI adapter barcodes varied by at least two base positions (Table SM1, SM2) which is necessary for an almost bias-free barcoding of individuals (Peterson, Weber, Kay, Fisher, & Hoekstra, 2012). For efficient multiplexing in Illumina sequencing, we used two Taq multiplex indices, 6 and 12, according to TruSeq guidelines for pooling with two-multiplexing (Illumina, TruSeq library prep pooling).

For creation of EcoRI and Taq adapters, corresponding complementary top and bottom P1.1 and P1.2 oligonucleotides (each 100 µM) were combined in a 1:1 ratio, together with annealing buffer of a final concentration of 1x (AB; 10x AB = 500 mM NaCl, 100 mM Tris/Cl pH 7.5). Annealing took place for 2.5 min at 98°C and following cooling down to room temperature at a rate of 2°C/min in a Labcycler (SensoQuest, Göttingen, Germany), resulting in a 40 µM solution per Eco- and Taq-site adapters. Final working concentration for Eco-site adapters was 0.5 µM, and for Taq-site adapters was 5 µM, using 1x AB for dilution. Taq-site top P2.2 adapters were biotinized at the 3' end.

We used four degenerated bases (equal mixture of A, C, G, T nucleotides at each nucleotide position) within the Taq-site adapter to distinguish polymerase chain reaction (PCR) duplicates during subsequent bioinformatic analysis (Table SM3; (Tin, Rheindt, Cros, & Mikheyev, 2015).

#### Digestion

Standardized DNA was used with a concentration of 120 – 250 ng/µl. For the first digestion, 26 µl standardized DNA with 0.5 µl EcoRI-HF NEB, and 3 µl 10x Smartcut buffer NEB was incubated for 45 min at 37°C. Following, 0.5 µl Taq$^{\alpha}$I NEB and 0.5 µl 1x Smartcut buffer NEB were added to the first digestion solution, incubating for 45 min at 65°C. All digestion steps were done in a Labcycler (SensoQuest, Göttingen, Germany).

#### Purification

DNA digest and custom-made solid-phase reversible immobilization (SPRI) bead solution (Jolivet & Foley, 2017) were mixed in a 1:1 ratio and incubated for 15 min at room temperature. Separation of beads with target DNA fragments and supernatant to be discarded was achieved using a magnetic stand. Keeping the reaction tube on the magnetic stand, tubes were cleaned twice with 70% ethanol, and dried for 10-15 min removing all ethanol. Dried beads were resuspended in 20 µl $H_2O$, with an incubation time of 2 min. Using a magnetic stand, purified DNA was transferred into ligation mix (see following step, Ligation).

Ligation

Shortly before needed and constantly cooled, a ligation mix for each Taq-site adapter was prepared, each consisting of: 2 µl Taq-site adapter (5 µM), 3 µl T4 10x ligase buffer, 1 µl T4 ligase (400U/µl; New England BioLabs Inc.), and 2 µl $H_2O$. To each ligation mix, 20 µl of purified DNA (see Purification), and 2 µl Eco-site adapters (0.5 µM) were added and incubated for 25 min at 23°C and 10 min at 65°C in a Labcycler (SensoQuest, Göttingen, Germany).

Pooling and size selection

All samples with the same Taq-site adapter but different Eco-site adapters were pooled together. From this pool, 300 µl (or at least 150 µl) were taken and AMPure XP beads (Beckman Coulter, Indianapolis, US) with a ratio of 0.7 were added. After incubating this mix for 10 min at room temperature, and separation of beads on a magnetic stand until supernatant was clear, the supernatant was saved. It was then mixed with AMPure XP beads (Beckman Coulter, Indianapolis, US) in a ratio of 0.12, incubated for 10 min at room temperature, and bead separation until clear supernatant on a magnetic stand was done. This time, however, the supernatant was discarded. Beads with target fragments were cleaned twice with 70% ethanol and dried for 10 min. Purified beads were eluted in 30 µl $H_2O$, incubated for 2 min, and separated on a magnetic stand. DNA concentration of purified size-selected library solution was measured (Qubit fluorometer; ThermoFisher Scientific, Waltham, MA, USA). The resulting libraries contained fragments with average length of 450 bp.

Selection for Taq-site biotin labelled adapters

Shortly before needed, 15 µl of M-270 Streptavidin Dynabeads (Dynal, Invitrogen, ThermoFisher Scientific, Waltham, MA, USA) were washed three times with 100 µl 1x binding and washing buffer (B&W buffer; 2x B&W buffer = 1 M Tris-HCl pH 8, 0.5 M EDTA, 5 M NaCl), and resuspended in 2x B&W buffer in twice the original volume (i.e. 30 µl). Size-selected DNA from previous step (see Pooling and size selection) was added in a 1:1 ratio and incubated for 15 min at room temperature with flicking the reaction tube every 5 min. For selection, Dynabeads were separated on a magnetic stand, and the supernatant was discarded. Subsequently, beads were washed again three times with 1x B&W buffer and resuspended in 45 µl $H_2O$.

PCR amplification and final clean-up

For PCR, the following ingredients were added to 45 µl of Dynabead suspension from the previous step (see Selection for Taq-site biotin labelled adapters): 3 µl primer 1 and 2 (each 10 µM; Table SM4) respectively, and 50 µl KAPA HiFi Hotstart Ready Mix (KAPA Biosystems, Wilmington, MA, USA). The resulting PCR suspension was divided into four tubes, to counteract competitive PCR bias, and ran on a LabCycler (SensoQuest, Göttingen, Germany) with the following conditions: initial denaturation for 2 min at 95°C, followed by 9 cycles of 20 s at 98°C, 30 s at 65°C, 30 s at 72°C. Following PCR, the four PCR solutions were separated on a magnetic stand, and supernatants were saved and combined. For final clean-up, to exclude PCR primers and short fragments, self-made SPRI bead solution in a ratio of 0.6 was added to PCR supernatant solution and cleaned as in the earlier purification step (see Purification). Purified libraries were eluted in 20 µl $H_2O$.

### Molarity of final ddRAD library and sequencing

The resulting ddRAD libraries were analysed for their DNA concentration, and molarity was calculated according to mean fragment size:

$$M[nM] = \frac{C}{F \cdot M_{bp}} \cdot 6, \quad (1)$$

where $M$ is the molarity of the final library given in nM, with a mean fragment length F (in bp; i.e. 450 bp), calculated with the overall library concentration $C$ (in ng/µl) after final purification and $M_{bp} = 660\ g/mol$.

Table SM1. Oligonucleotide sequences of top and bottom part of EcoRI adapters. EcoRI adapter motif is highlighted in bold (see all 48 motifs used in this study in Table SM2).

| adapter part | EcoRI adapter sequence (5' – 3') |
|---|---|
| top (P1.2) | ACACTCTTTCCCTACACGACGCTCTTCCGATCTTCGAT |
| bottom (P1.1) | **AATTA**TCGAAGATCGGAAGAGCGTCGTGTAGGGAAAGAGTGT |

Table SM2. Oligonucleotide motif of 48 EcoRI adapters used in the study.

| # | EcoRI motif | # | Eco-site adapter nucleotide sequence |
|---|---|---|---|
| 01 | GCATG | 25 | CTGCG |
| 02 | AACCA | 26 | CTGTC |
| 03 | CGATC | 27 | CTTGG |
| 04 | TCGAT | 28 | GACAC |
| 05 | TGCAT | 29 | GAGAT |
| 06 | CAACC | 30 | GAGTC |
| 07 | GGTTG | 31 | GCCGT |
| 08 | AAGGA | 32 | GCTGA |
| 09 | AGCTA | 33 | GGATA |
| 10 | ACACA | 34 | GGCCA |
| 11 | AATTA | 35 | GGCTC |
| 12 | ACGGT | 36 | GTAGT |
| 13 | ACTGG | 37 | GTCCG |
| 14 | ACTTC | 38 | GTCGA |
| 15 | ATACG | 39 | TACCG |
| 16 | ATGAG | 40 | TACGT |
| 17 | ATTAC | 41 | TAGTA |
| 18 | CATAT | 42 | TATAC |
| 19 | CGAAT | 43 | TCACG |

| | | | |
|---|---|---|---|
| 20 | CGGCT | 44 | TCAGT |
| 21 | CGGTA | 45 | TCCGG |
| 22 | CGTAC | 46 | TCTGC |
| 23 | CGTCG | 47 | TGGAA |
| 24 | CTGAT | 48 | TTACC |

Table SM3. Oligonucleotide sequences of top and bottom part of Taq$^{\alpha}$I adapters. Top part of Taq adapter is phosphorylated at 5'-end and biotin labelled at 3'-end. Random bases are indicated with *N*. Bold nucleotides in Taq adapter sequences indicate Taq motif.

| Taq index | adapter part | Taq motif | Taq adapter sequence (5' – 3') |
|---|---|---|---|
| both | top (P2.2) | - | [Phos]CGGATCGGAAGAGCACACGTCTGAACTCCAGT CAC[BtnTg] |
| 6 | bottom (P2.1) | GCCAAT | CAAGCAGAAGACGGCATACGAGAT*NNNN***ATTGGC**GTGAC TGGAGTTCAGACGTGTGC |
| 12 | Bottom (P2.1) | CTTGTA | CAAGCAGAAGACGGCATACGAGAT*NNNN***TACAAG**GTGAC TGGAGTTCAGACGTGTGC |

Table SM4. PCR primers.

| PCR primer | PCR primer sequence (5' – 3') |
|---|---|
| 1 | AATGATACGGCGACCACCGAGATCTACACTCTTTCCCTACACGACG |
| 2 | CAAGCAGAAGACGGCATACGA |

### Supplemental Figures

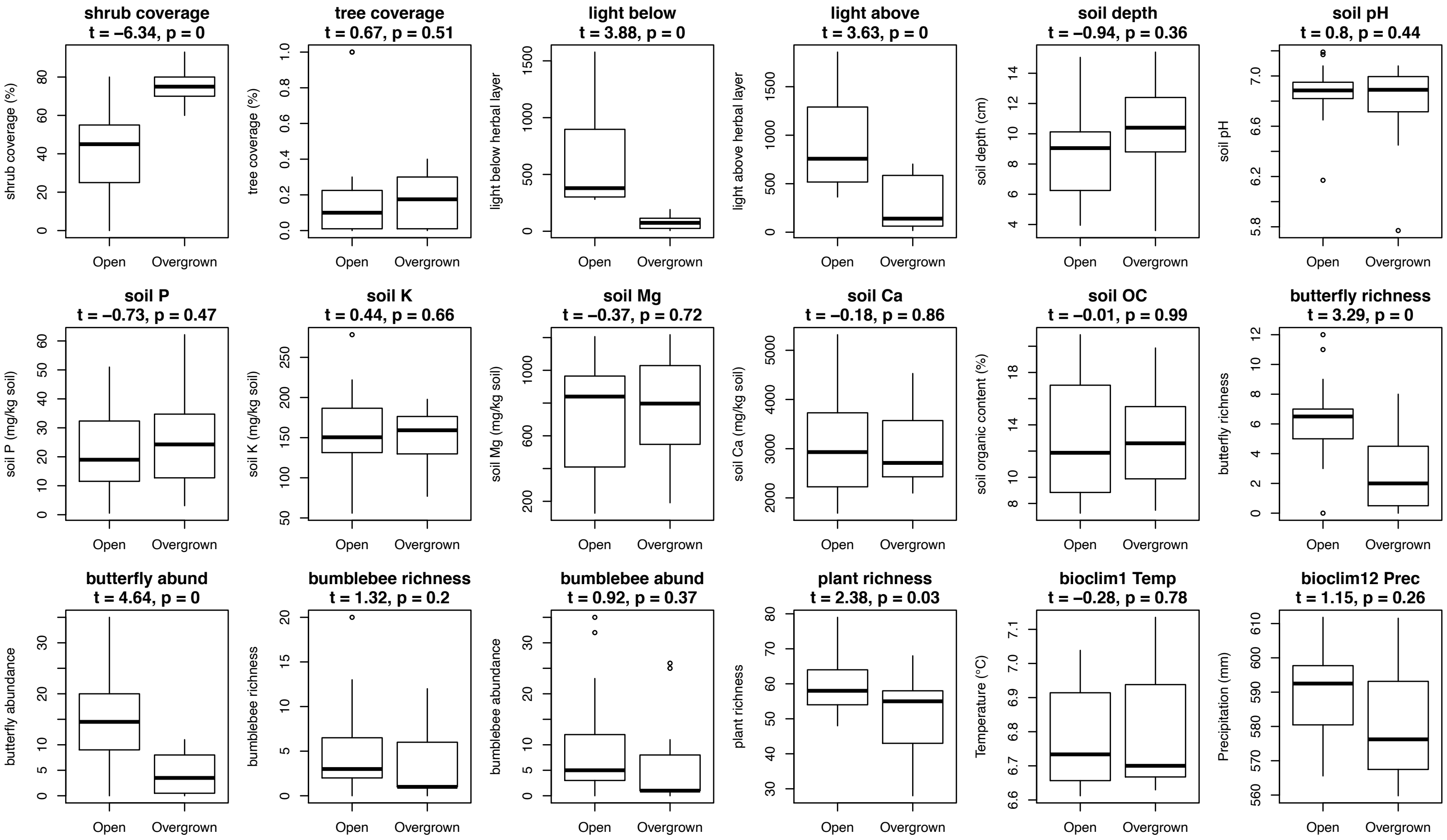

**Figure S1** Boxplot of environmental variables and their response to habitat type (open – overgrown) for *Primula veris* populations on the islands Muhu and Saaremaa, Estonia. In the title of each panel, the name of the respective environmental variable is indicated, together with t-statistics from a (non-paired) two-sample t-test.

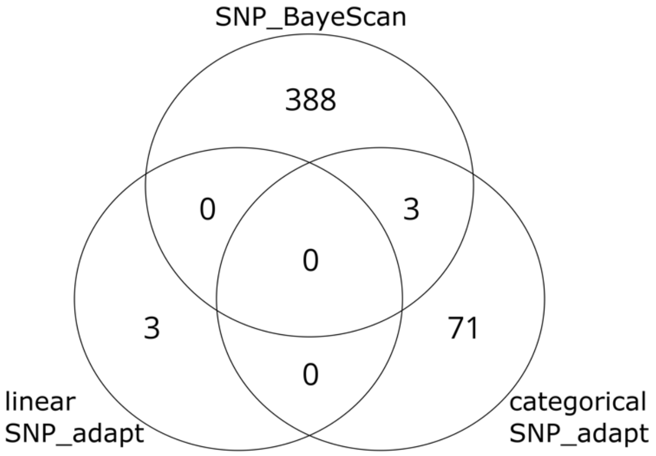

**Figure S2** Venn diagram of shared putatively adaptive loci of *Primula veris* identified by different methods: linear SNP_adapt – association between SNPs and environmental variables (which are significantly different in the open and overgrown habitats) resulting from latent factor mixed effect model analysis (LFMM2); categorical SNP_adapt – SNPs significant in three paired tests (paired t-test, paired Wilcoxon signed-rank test, and sign test), using the 20 populations that were sampled in pairs per study site; SNP_BayeScan – SNPs under potential diversifying or balancing selection, resulting from $F_{ST}$ outlier tests.

**Figure S3** Please see separate file.

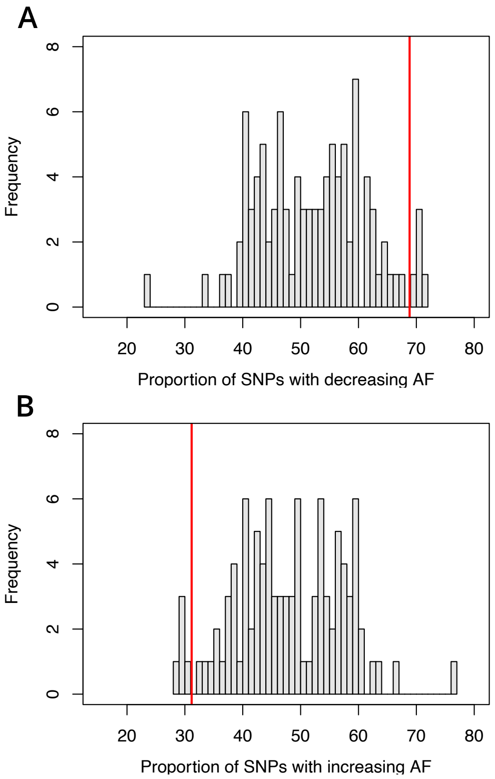

**Figure S4** Distribution of the frequency of the proportion of SNPs with decreasing (A) or increasing (B) beneficial allele frequency (AF) from open to overgrown habitats from SNP_adapt of 100 randomised data sets. The red line indicates the value of the proportion of SNPs with decreasing or increasing beneficial AF from SNP_adapt of the original data set. The proportion was calculated as the respective number of SNPs with de- or increasing beneficial AF divided by the total number of putatively adaptive SNPs for the respective data set (original or randomised).

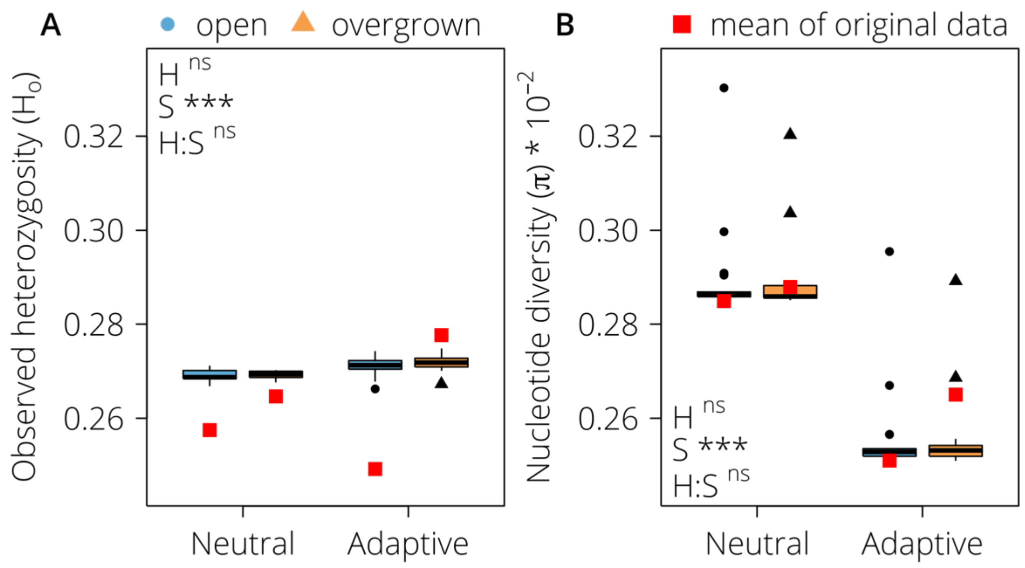

**Figure S5** Boxplots of observed heterozygosity ($H_o$; A) and nucleotide diversity ($\pi$; B) at putatively neutral and adaptive loci in open (blue, circles) and overgrown (orange, triangles) grasslands. $H_o$ and $\pi$ values are averaged per population over 100 randomised data sets. Red squares indicate the mean of $H_o$ or $\pi$ of the original data of the respective habitat – SNP set combination. Significance levels are given for the model testing using randomised data. Factors: habitat (H): open and overgrown; SNP_set (S): SNP_neutral and SNP_adaptive. Significance values: $^{ns}$ $p > 0.05$; *** $p < 0.001$.

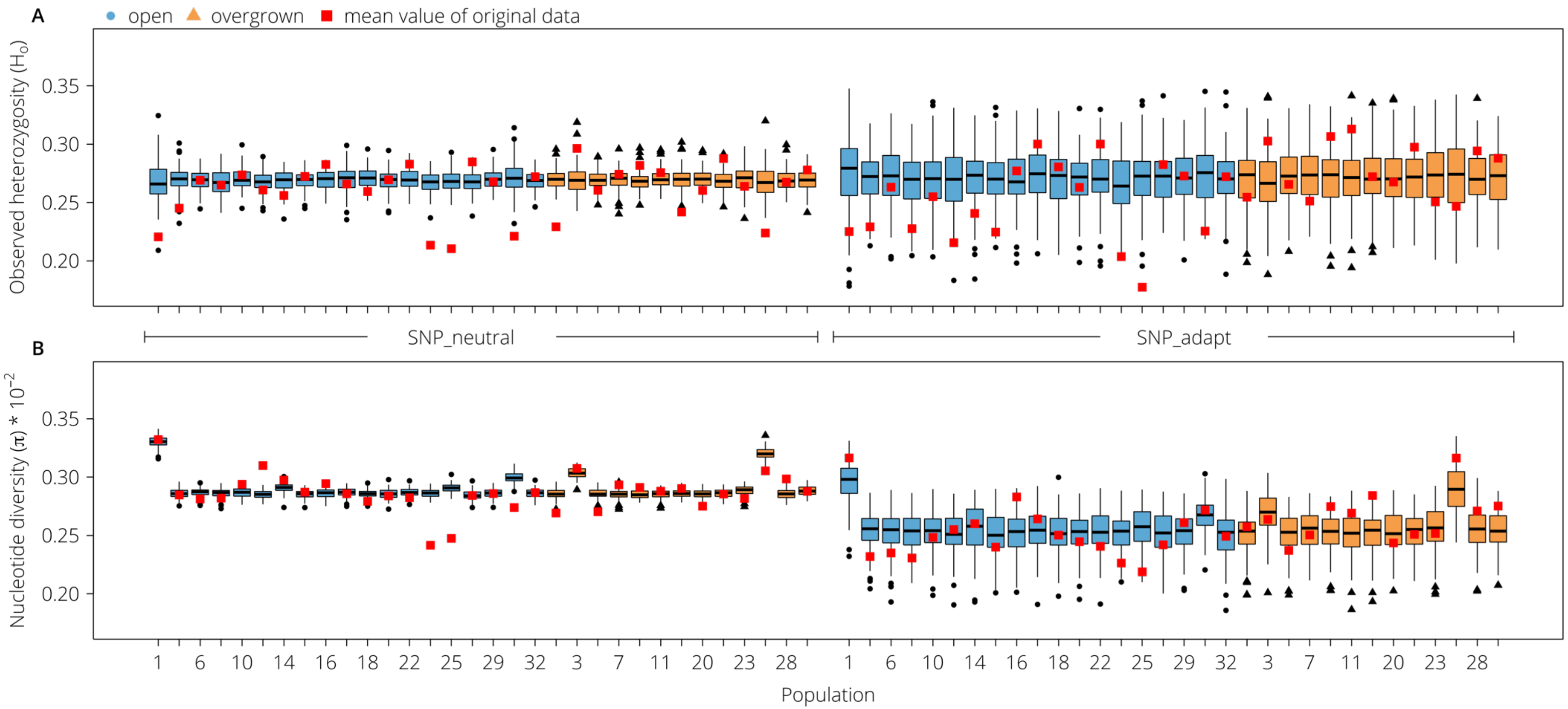

**Figure S6** Boxplots of genetic diversity (observed heterozygosity, $H_o$; A) and nucleotide diversity ($\pi$; B) of *Primula veris* per population of 100 randomised data sets, using the respective putatively neutral and and adaptive SNP sets in open and overgrown grasslands. Red squares indicate the population mean of $H_o$ or $\pi$ of the respective population. Population numbers and associations to habitat types can be found in Table 1 in the main manuscript.

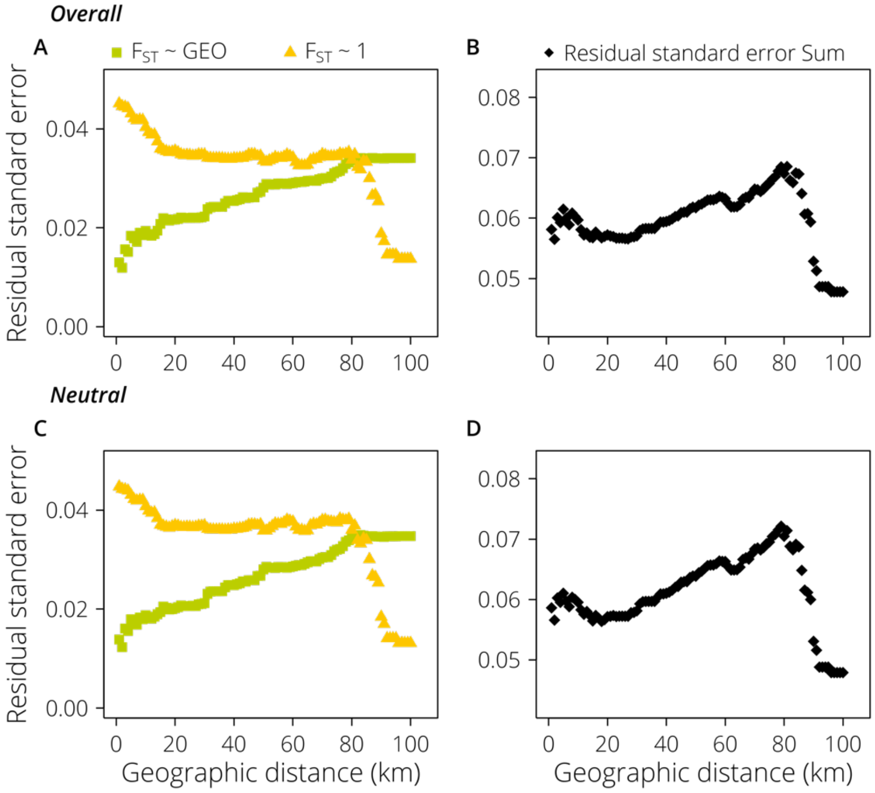

**Figure S7** Residual standard errors for (A, C) a set of linear models with $F_{ST}$ ~ geographic distance (GEO) between 1 and 100 km distance between population pairs (green squares), and a complementary set of linear models with $F_{ST}$ ~ 1 (yellow triangles), measured using the overall (3,084 loci) and neutral (2,619 loci) SNP set, respectively. In (B, D) the sum of residual standard errors of both complementary models at a specific geographic distance is presented, measured using the overall (3,084 loci) and neutral (2,619 loci) SNP set, respectively.

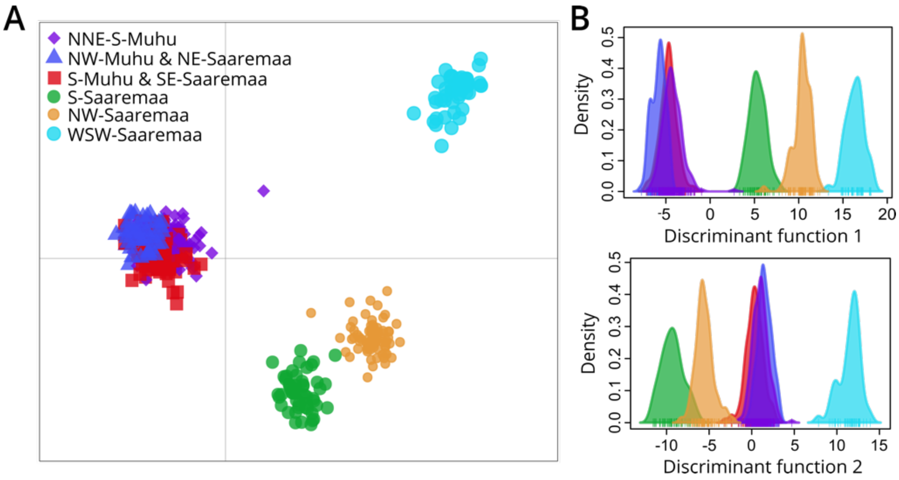

**Figure S8** Genetic structure of *Primula veris* populations in the study region. (A) Result of the discriminant analysis of principle components (DAPC) on the overall (3,084 loci) SNP set, respectively, resulting in six geographical clusters (named by cardinal directions). The distribution of the six clusters per discriminant function is shown in panels B.

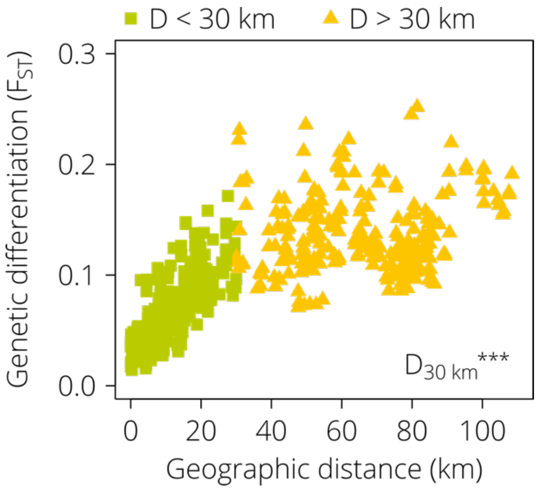

**Figure S9** Relationship between genetic differentiation ($F_{ST}$) and geographic distance for all possible pairs of 32 *Primula veris* populations, measured at the overall (3,084 loci) SNP set. Population pairs with a geographic distance (D) less or equal to 30 km are visualized as green squares. Population pairs with a geographic distance greater than 30 km are given as yellow triangles. The effect of geographic distance on $F_{ST}$ is highly significant ($p < 0.001$) up to a threshold of 30 km.

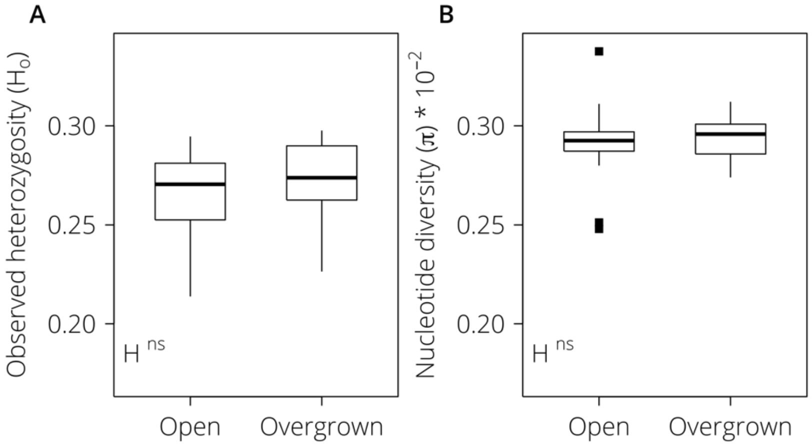

**Figure S10** Boxplots of genetic diversity (observed heterozygosity, $H_o$; A) and nucleotide diversity ($\pi$; B) of *Primula veris* using the overall SNP set (3,084 loci) in open and overgrown historical grasslands. Factor: habitat (H): open and overgrown. Significance values: $^{ns}$ $p > 0.05$.

Supplemental Tables

**Table S1** Number of associations between SNPs and environmental variables in the latent factor mixed effect model (LFMM2) analysis. Environmental variables significantly (t-test, $p \leq 0.05$) different in open and overgrown habitats, and the corresponding amount of SNP associations are indicated in bold.

| Variable | Significance (*p*) between open - overgrown | Number of associations (LFMM) |
|---|---|---|
| Shrub coverage | **< 0.001** | **0** |
| Tree coverage | 0.51 | 0 |
| Light below herbal layer | **< 0.01** | **0** |
| Light above herbal layer | **< 0.01** | **0** |
| Soil depth | 0.36 | 0 |
| Soil pH | 0.44 | 2 |
| Soil P | 0.47 | 0 |
| Soil K | 0.66 | 2 |
| Soil Mg | 0.72 | 0 |
| soil Ca | 0.86 | 0 |
| Soil organic content | 0.99 | 0 |
| Butterfly richness | **< 0.01** | **0** |
| Butterfly abundance | **< 0.001** | **3** |
| Bumblebee richness | 0.20 | 1 |
| Bumblebee abundance | 0.37 | 0 |
| Plant richness | **0.03** | **0** |
| Bioclim1 (temperature) | 0.78 | 0 |
| Bioclim12 (precipitation) | 0.26 | 0 |

**Table S2** Results of best-fit generalized mixed effect models for the effect of geographic distance (D) and habitat type distance ($H_D$ – open, overgrown, or between habitats) on pairwise genetic differentiation ($F_{ST}$) of *Primula veris* populations, for different SNP sets (SNP_neutral, SNP_overall). l-95% and u-95% – lower or upper limit of 95% confidence interval, respectively. DIC differences ($DIC_{diff}$) between the full model ($F_{ST}$ ~ D + $H_D$) and the simplified model to test for the importance of each effect is given (importance of D: $F_{ST}$ ~ $H_D$; importance of $H_D$: $F_{ST}$ ~ D). Model results for each SNP set with the best fit according to model selection with DIC are presented.

| Effect | Effect size | l-95% | u-95% | $p_{MCMC}$ | $DIC_{diff}$ |
|---|---|---|---|---|---|
| **SNP_neutral** | | | | | |
| D | 8.0e$^{-04}$ | 7.0e$^{-04}$ | 9.0e$^{-04}$ | < 0.001 | -189.48 |
| $H_D$ | - | - | - | - | 5.83 |
| ***SNP_overall*** | | | | | |
| D | 8.5e$^{-04}$ | 7.6e$^{-04}$ | 9.6e$^{-04}$ | < 0.001 | -219.56 |
| $H_D$ | - | - | - | - | 5.73 |

**Table S3** Genetic diversity measures using the overall set of loci (3,084 SNPs) for the studied populations of *Primula veris*. Habitat – affiliation to open or overgrown habitat; pair – affiliation to a pair of closely situated populations (0 indicates affiliation to no pair); $H_o$ – observed heterozygosity; $\pi$ – nucleotide diversity.

| | | | | Overall SNP set | |
|---|---|---|---|---|---|
| **Pop ID** | **Region** | **Habitat** | **Pair** | $H_o$ | $\pi$ |
| **1** | Saaremaa | open | 1 | 0.23 | 0.0034 |
| **2** | Saaremaa | overgrown | 1 | 0.24 | 0.0027 |
| **3** | Saaremaa | overgrown | 0 | 0.30 | 0.0031 |
| **4** | Saaremaa | open | 0 | 0.25 | 0.0029 |
| **5** | Muhu | overgrown | 2 | 0.27 | 0.0028 |
| **6** | Muhu | open | 2 | 0.29 | 0.0029 |
| **7** | Saaremaa | overgrown | 3 | 0.27 | 0.0030 |
| **8** | Saaremaa | open | 3 | 0.26 | 0.0029 |
| **9** | Muhu | overgrown | 4 | 0.29 | 0.0030 |
| **10** | Muhu | open | 4 | 0.28 | 0.0030 |
| **11** | Muhu | overgrown | 0 | 0.28 | 0.0030 |
| **12** | Saaremaa | open | 0 | 0.25 | 0.0031 |
| **13** | Muhu | overgrown | 5 | 0.25 | 0.0029 |
| **14** | Muhu | open | 5 | 0.26 | 0.0030 |
| **15** | Saaremaa | open | 0 | 0.27 | 0.0029 |
| **16** | Saaremaa | open | 0 | 0.28 | 0.0030 |
| **17** | Muhu | open | 0 | 0.28 | 0.0029 |
| **18** | Muhu | open | 0 | 0.27 | 0.0029 |
| **19** | Saaremaa | open | 6 | 0.28 | 0.0029 |
| **20** | Saaremaa | overgrown | 6 | 0.27 | 0.0029 |
| **21** | Muhu | overgrown | 7 | 0.30 | 0.0029 |
| **22** | Muhu | open | 7 | 0.29 | 0.0029 |
| **23** | Saaremaa | overgrown | 0 | 0.26 | 0.0028 |
| **24** | Saaremaa | open | 0 | 0.22 | 0.0025 |
| **25** | Saaremaa | open | 8 | 0.21 | 0.0025 |
| **26** | Saaremaa | overgrown | 8 | 0.23 | 0.0031 |
| **27** | Muhu | open | 0 | 0.29 | 0.0029 |
| **28** | Muhu | overgrown | 9 | 0.28 | 0.0030 |
| **29** | Muhu | open | 9 | 0.27 | 0.0029 |
| **30** | Saaremaa | open | 0 | 0.22 | 0.0028 |
| **31** | Muhu | overgrown | 10 | 0.29 | 0.0030 |
| **32** | Muhu | open | 10 | 0.28 | 0.0029 |
| **Open mean** | | | | 0.26 | 0.0029 |
| **Overgrown mean** | | | | 0.27 | 0.0030 |

**Table S4** Results of linear mixed effect models for observed heterozygosity ($H_o$) and nucleotide diversity ($\pi$), for the overall SNP set (SNP_overall) compiling 3,084 loci. Fitted parameters (± SE), t-values and significance ($p_{Model}$) are given for the full model of each genetic measure, $H_o$ and $\pi$. Factors: habitat (H): open and overgrown. $RF_{region}$ denotes the use of region (Muhu and Saaremaa) as random factor (RF). The importance of habitat is given as $\chi^2$ and p-values for each genetic measure, tested by the comparison of the full model with the simplest model (no interaction or fixed factor).

| Effect | Value | SE | t | $p_{Model}$ | $\chi^2$ | *p* |
|---|---|---|---|---|---|---|
| **$H_o$ ~ H + $RF_{region}$** | | | | | **Importance of H** | |
| habitat | 0.007 | 0.007 | 0.901 | 0.375 | 0.860 | 0.354 |
| **$\pi$ ~ H + $RF_{region}$** | | | | | ***Importance of H*** | |
| habitat | $3.6e^{-5}$ | $5.9e^{-5}$ | 0.603 | 0.551 | 0.386 | 0.535 |

### References

Jolivet, P., & Foley, J. (2017). SPRI bead mix preparation protocol. https://openwetware.org/wiki/SPRI_bead_mix

Peterson, B. K., Weber, J. N., Kay, E. H., Fisher, H. S., & Hoekstra, H. E. (2012). Double digest RADseq: an inexpensive method for de novo SNP discovery and genotyping in model and non-model species. *PLoS ONE*, *7*. doi: 10.1371/journal.pone.0037135

Tin, M. M. Y., Rheindt, F. E., Cros, E., & Mikheyev, A. S. (2015). Degenerate adaptor sequences for detecting PCR duplicates in reduced representation sequencing data improve genotype calling accuracy. *Molecular Ecology Resources*, *15*, 329–336. doi: 10.1111/1755-0998.12314

Lepais, O., & Weir, J. T. (2014). SimRAD: an R package for simulation-based prediction of the number of loci expected in RADseq and similar genotyping by sequencing approaches. *Molecular Ecology Resources*, *14*, 1314–1321. doi: 10.1111/1755-0998.12273
