## Supporting Information Figure S3 for "The effect of recent habitat change on genetic diversity at putatively adaptive and neutral loci in *Primula veris* in semi-natural grasslands"

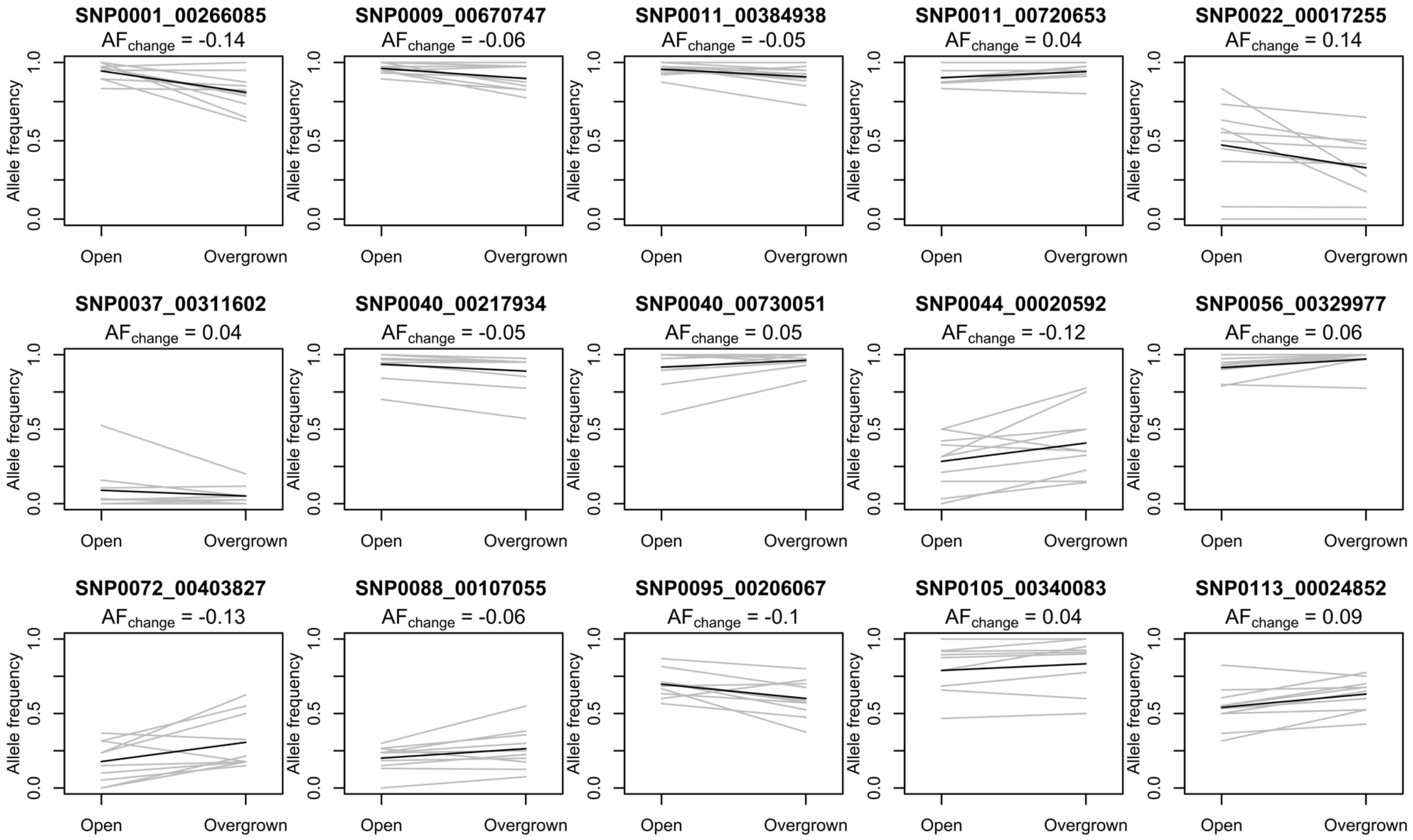
SNP0001_00266085
AF$_{change}$ = -0.14
SNP0009_00670747
AF$_{change}$ = -0.06
SNP0011_00384938
AF$_{change}$ = -0.05
SNP0011_00720653
AF$_{change}$ = 0.04
SNP0022_00017255
AF$_{change}$ = 0.14
SNP0037_00311602
AF$_{change}$ = 0.04
SNP0040_00217934
AF$_{change}$ = -0.05
SNP0040_00730051
AF$_{change}$ = 0.05
SNP0044_00020592
AF$_{change}$ = -0.12
SNP0056_00329977
AF$_{change}$ = 0.06
SNP0072_00403827
AF$_{change}$ = -0.13
SNP0088_00107055
AF$_{change}$ = -0.06
SNP0095_00206067
AF$_{change}$ = -0.1
SNP0105_00340083
AF$_{change}$ = 0.04
SNP0113_00024852
AF$_{change}$ = 0.09
Allele frequency
0.0
0.5
1.0
Open
Overgrown

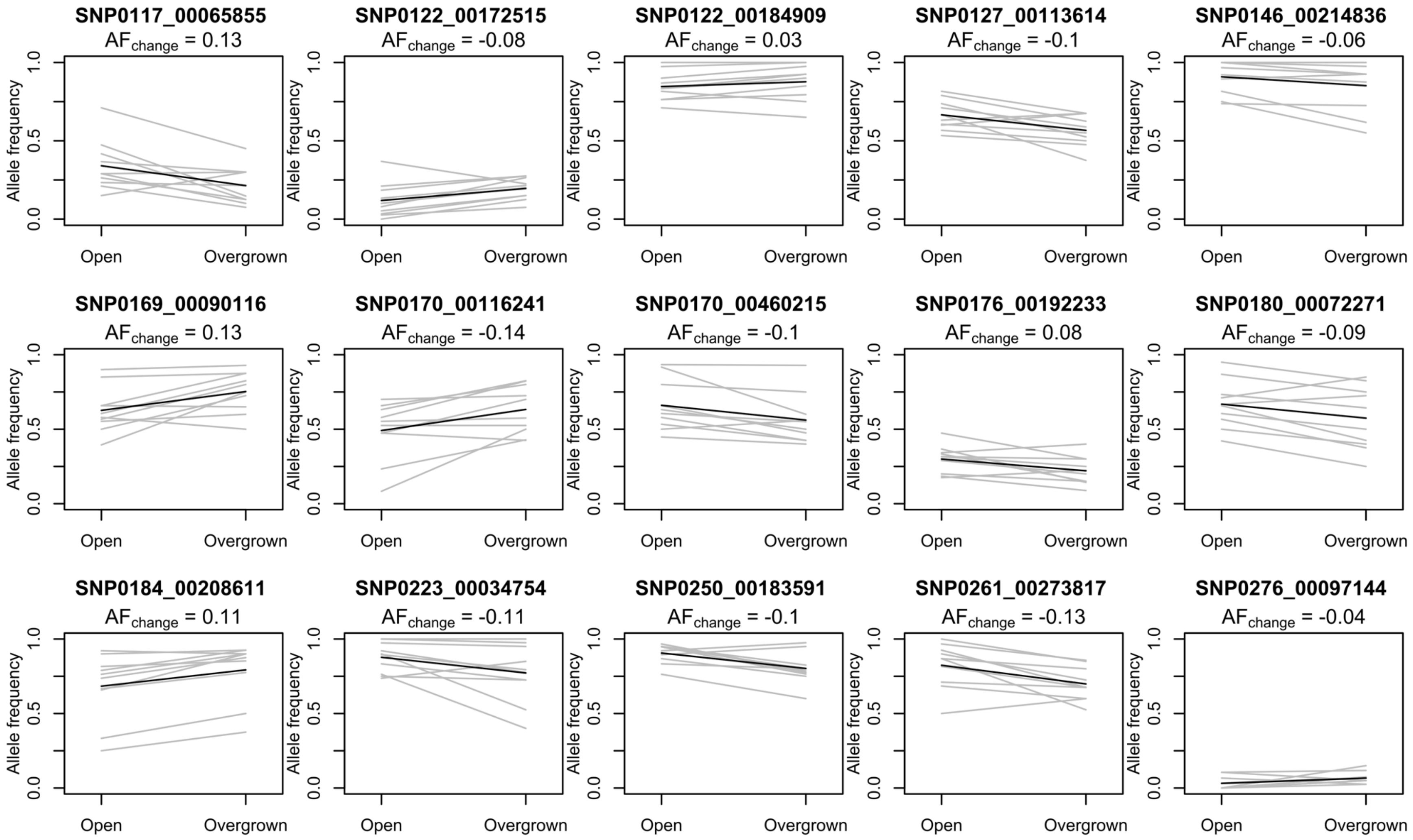

SNP0117_00065855
$AF_{change}$ = 0.13
SNP0122_00172515
$AF_{change}$ = -0.08
SNP0122_00184909
$AF_{change}$ = 0.03
SNP0127_00113614
$AF_{change}$ = -0.1
SNP0146_00214836
$AF_{change}$ = -0.06
SNP0169_00090116
$AF_{change}$ = 0.13
SNP0170_00116241
$AF_{change}$ = -0.14
SNP0170_00460215
$AF_{change}$ = -0.1
SNP0176_00192233
$AF_{change}$ = 0.08
SNP0180_00072271
$AF_{change}$ = -0.09
SNP0184_00208611
$AF_{change}$ = 0.11
SNP0223_00034754
$AF_{change}$ = -0.11
SNP0250_00183591
$AF_{change}$ = -0.1
SNP0261_00273817
$AF_{change}$ = -0.13
SNP0276_00097144
$AF_{change}$ = -0.04
Allele frequency
0.0
0.5
1.0
Open
Overgrown

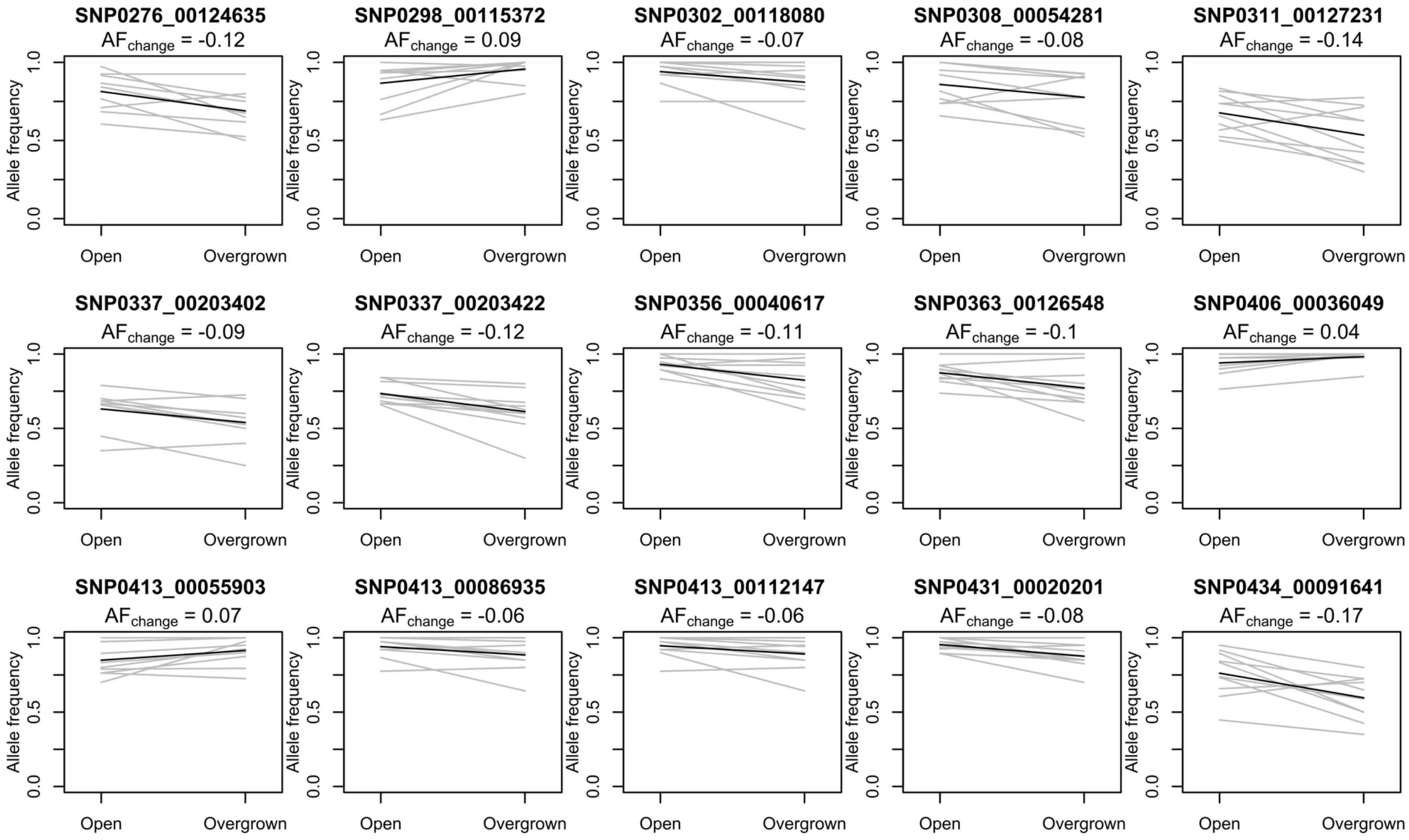

SNP0276_00124635
AFchange = -0.12
SNP0298_00115372
AFchange = 0.09
SNP0302_00118080
AFchange = -0.07
SNP0308_00054281
AFchange = -0.08
SNP0311_00127231
AFchange = -0.14
SNP0337_00203402
AFchange = -0.09
SNP0337_00203422
AFchange = -0.12
SNP0356_00040617
AFchange = -0.11
SNP0363_00126548
AFchange = -0.1
SNP0406_00036049
AFchange = 0.04
SNP0413_00055903
AFchange = 0.07
SNP0413_00086935
AFchange = -0.06
SNP0413_00112147
AFchange = -0.06
SNP0431_00020201
AFchange = -0.08
SNP0434_00091641
AFchange = -0.17
Allele frequency
0.0
0.5
1.0
Open
Overgrown

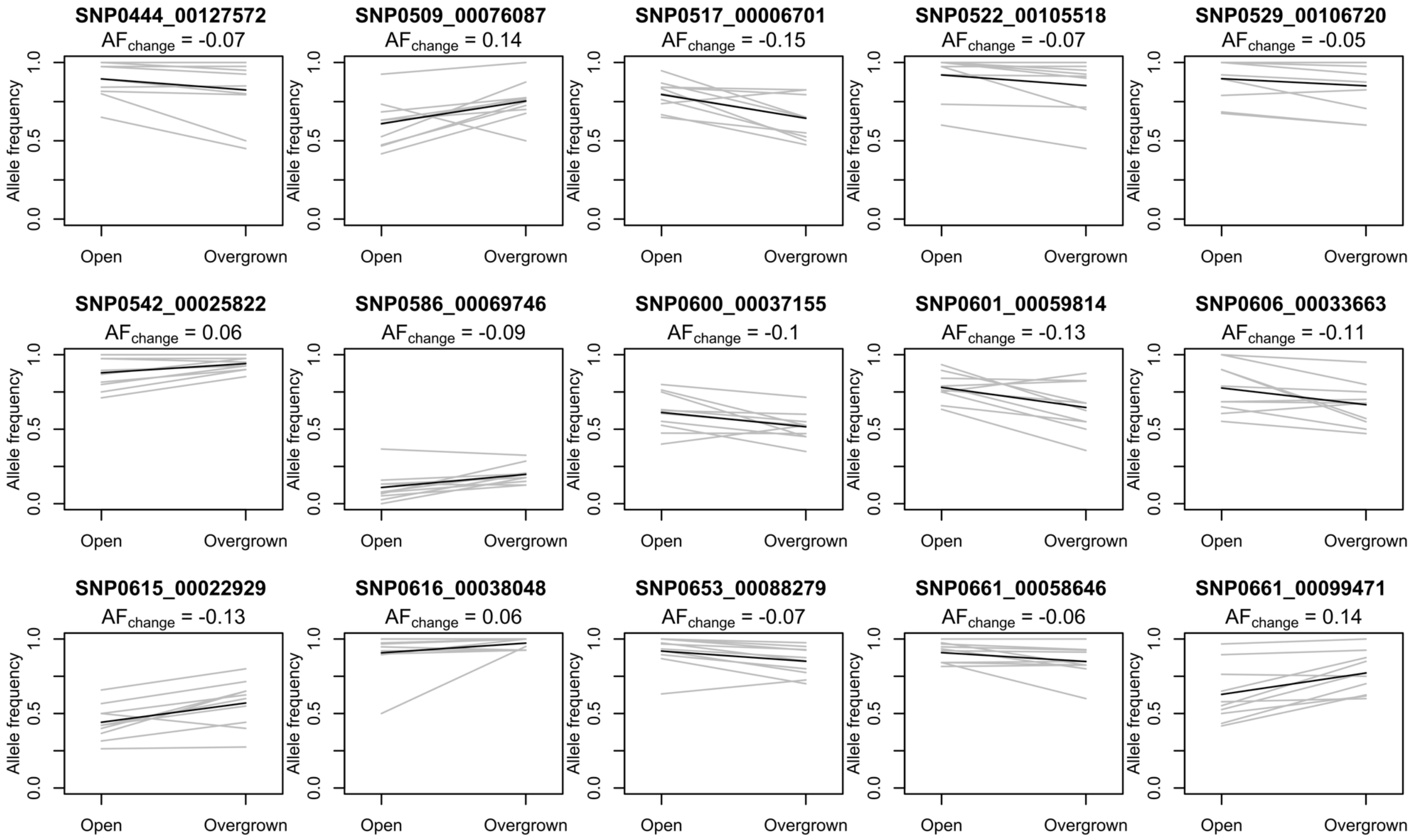

SNP0444_00127572
AF change = -0.07
SNP0509_00076087
AF change = 0.14
SNP0517_00006701
AF change = -0.15
SNP0522_00105518
AF change = -0.07
SNP0529_00106720
AF change = -0.05
SNP0542_00025822
AF change = 0.06
SNP0586_00069746
AF change = -0.09
SNP0600_00037155
AF change = -0.1
SNP0601_00059814
AF change = -0.13
SNP0606_00033663
AF change = -0.11
SNP0615_00022929
AF change = -0.13
SNP0616_00038048
AF change = 0.06
SNP0653_00088279
AF change = -0.07
SNP0661_00058646
AF change = -0.06
SNP0661_00099471
AF change = 0.14
Allele frequency
0.0
0.5
1.0
Open
Overgrown

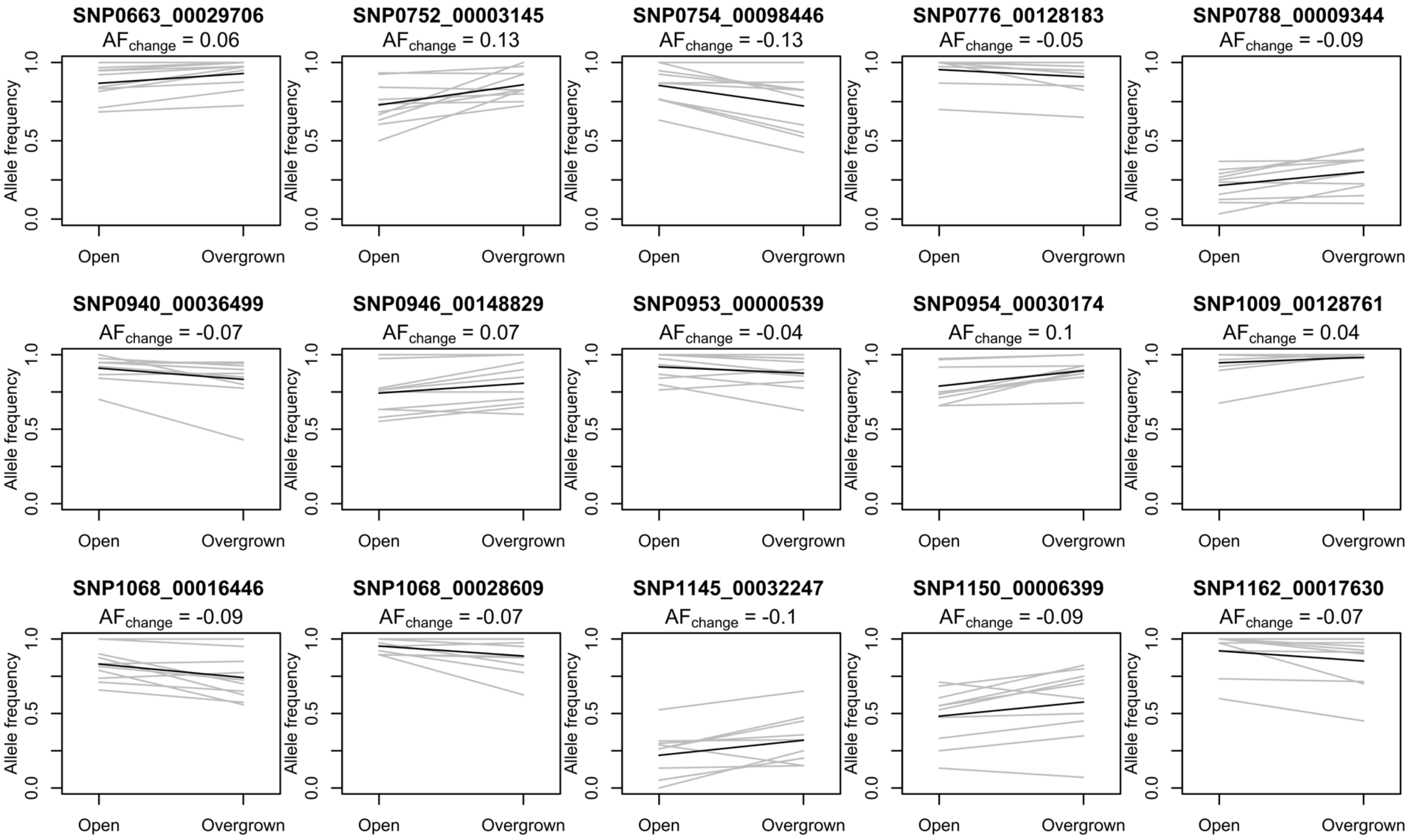
SNP0663_00029706
AFchange = 0.06
SNP0752_00003145
AFchange = 0.13
SNP0754_00098446
AFchange = -0.13
SNP0776_00128183
AFchange = -0.05
SNP0788_00009344
AFchange = -0.09
SNP0940_00036499
AFchange = -0.07
SNP0946_00148829
AFchange = 0.07
SNP0953_00000539
AFchange = -0.04
SNP0954_00030174
AFchange = 0.1
SNP1009_00128761
AFchange = 0.04
SNP1068_00016446
AFchange = -0.09
SNP1068_00028609
AFchange = -0.07
SNP1145_00032247
AFchange = -0.1
SNP1150_00006399
AFchange = -0.09
SNP1162_00017630
AFchange = -0.07
Allele frequency
0.0
0.5
1.0
Open
Overgrown

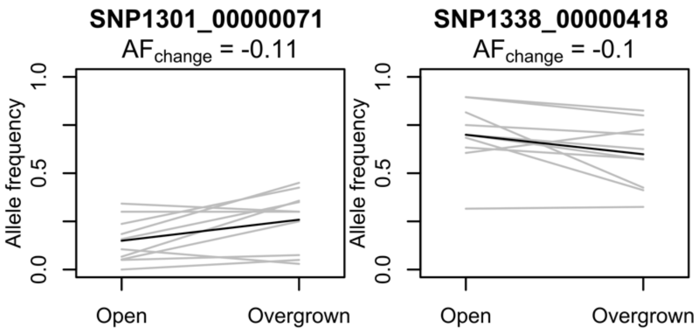


**Figure S3** Patterns of allele frequencies (AF) between populations pairs (from open and overgrown habitats) for the 77 putatively adaptive SNPs derived from linear and categorical environmental association analyses (SNP_adapt). In the title of each panel, the locus name and the average frequency change (black line) of the beneficial allele from the open habitat towards the overgrown habitat is indicated. Grey lines depict the patterns in each population pair. Negative values indicate a lower beneficial allele frequency (for the open habitat) in the overgrown population compared to the open population, positive values indicate a higher beneficial allele frequency (for the open habitat) in the overgrown population compared to the open population.
